# *C9orf72* arginine-rich dipeptide repeat proteins disrupt importin β-mediated nuclear import

**DOI:** 10.1101/787473

**Authors:** Lindsey R. Hayes, Lauren Duan, Kelly Bowen, Petr Kalab, Jeffrey D. Rothstein

## Abstract

Disruption of nucleocytoplasmic transport (NCT), including mislocalization of the importin β cargo, TDP-43, is a hallmark of amyotrophic lateral sclerosis (ALS), including ALS caused by a hexanucleotide repeat expansion in *C9orf72*. However, the mechanism(s) remain unclear. Importin β and its cargo adaptors have been shown to co-precipitate with the *C9orf72*-arginine-containing dipeptide repeat proteins (R-DPRs), poly-glycine arginine (GR) and poly-proline arginine (PR), and are protective in genetic modifier screens. Here, we show that R-DPRs interact with importin β, disrupt its cargo loading, and inhibit nuclear import in permeabilized mouse neurons and HeLa cells, in a manner that can be rescued by RNA. Although R-DPRs induce widespread protein aggregation in this *in vitro* system, transport disruption is not due to NCT protein sequestration, nor blockade of the phenylalanine-glycine (FG)-rich nuclear pore complex. Our results support a model in which R-DPRs interfere with nuclear transport receptors in the vicinity of the nuclear envelope.

## Introduction

A GGGGCC hexanucleotide repeat expansion (HRE) in *C9orf72* is the most common known cause of amyotrophic lateral sclerosis (ALS) and is also a major cause of frontotemporal dementia (FTD) and the ALS /FTD overlap syndrome (DeJesus-Hernandez et al., 2011; Renton et al., 2011; Majounie et al., 2012). The *C9orf72* HRE is thought to cause disease by a toxic gain of function involving expanded repeat RNA and dipeptide repeat proteins (DPRs) produced by repeat-associated (non-AUG) translation, although a modest reduction in C9ORF72 protein is also seen (reviewed by Cook and Petrucelli, 2019). Predicted products of *C9orf72* HRE translation in both the sense (poly-GP, poly-GA, poly-GR) and antisense (poly-GP, poly-PR, poly-PA) directions have been identified in postmortem tissue (Zu et al., 2013; Ash et al., 2013; Mackenzie et al., 2013), and overexpression of a subset of DPRs, including poly-GA and the arginine-containing DPRs poly-GR and poly-PR (R-DPRs), is toxic in cell culture (May et al., 2014; Wen et al., 2014) and animal models (Zhang et al., 2016; 2018; 2019).

Growing evidence suggests that disruption of nucleocytoplasmic transport (NCT), the regulated trafficking of proteins and ribonucleoprotein complexes between the nucleus and cytoplasm, is a major pathophysiologic mechanism in neurodegenerative diseases (reviewed by Hutten and Dormann, 2019). Bidirectional NCT across the nuclear envelope occurs through nuclear pore complexes (NPC), which are large (125 MDa) assemblies comprised of multiple copies of ∼30 different nucleoporins (Nups) (Reichelt et al., 1990). Whereas small cargoes passively equilibrate across the NPC, larger cargoes are increasingly excluded by a matrix of natively-unfolded phenylalanine-glycine (FG)-rich nucleoporins lining the central channel (Timney et al., 2016; Frey et al., 2018; Mohr et al., 2009). Transport of restricted cargoes requires nuclear transport receptors (importins, exportins, and transportins, aka NTRs or karyopherins), which mediate the rapid transport of cargo through the FG-barrier (reviewed by (Pemberton and Paschal, 2005). The small GTPase Ran dictates the directionality of transport via a steep concentration gradient of RanGTP across the nuclear membrane, established by the nuclear guanine nucleotide exchange factor RCC1 and the cytoplasmic GTPase-activating protein RanGAP1. Nuclear RanGTP promotes importin-cargo unloading and exportin-cargo complex assembly, while the cytoplasmic conversion of RanGTP to RanGDP disassembles exportin-cargo complexes and enables importin-cargo binding. We and others have found evidence of NCT disruption in postmortem tissue and animal models of *C9orf72*-ALS, Alzheimer’s disease, and Huntington’s disease, including mislocalization and loss of Nups and disruption of the Ran gradient (Zhang et al., 2015; Grima et al., 2017; Eftekharzadeh et al., 2018).

Cytoplasmic mislocalization of the importin β cargo TDP-43, a predominantly nuclear DNA/RNA-binding protein that undergoes nucleocytoplasmic shuttling (Pinarbasi et al., 2018), is a major pathological hallmark of ALS, including *C9orf72*-ALS (Neumann et al., 2006; Mackenzie et al., 2014). Although importin β directly imports a subset of cargoes, most (including TDP-43) are recruited via a heterodimer with importin α, bound by its importin β-binding domain (IBB), leading to formation of a trimeric import complex (cargo•importin α•importin β) (reviewed by (Lott and Cingolani, 2011). Classical and non-classical nuclear localization signals (NLS) of importin β cargoes and the IBB (a disordered region that is also a functional NLS), are enriched in arginine and lysine residues that mediate high-affinity interactions within the import complex. *C9orf72* genetic modifier screens have identified a beneficial role for NTRs, including importin β and its importin α family of cargo adaptors (Zhang et al., 2015; Freibaum et al., 2015; Jovičić et al., 2015; Boeynaems et al., 2016; Kramer et al., 2018). Moreover, multiple interactome screens have shown that R-DPRs co-precipitate with importin β, indicating a possible direct interaction (Lee et al., 2016; Lin et al., 2016; Yin et al., 2017). We hypothesized that R-DPRs may directly interact with importin β by mimicking the arginine-and lysine-rich IBB, disrupting nuclear import complex formation.

Here, we use FRET and biochemical assays to show that R-DPRs interact with importin β, disrupt import complex formation, and confer dose- and length-dependent disruption of importin β-mediated nuclear import in the permeabilized cell assay, which we adapted for primary neurons. Addition of R-DPRs to the transport assay triggers rapid formation of insoluble aggregates, which recruit numerous RNA-binding and ribosomal proteins, as well as NPC and NCT proteins. However, by separating the soluble and insoluble phases of the reaction, we show that transport disruption is not due to sequestration of NCT components, nor the ability of R-DPRs to impede passage through the NPC, but due to perturbation of soluble transport factors, an effect that is reversible by RNA. Taken together, these data support a model of R-DPR-based NCT inhibition via disruption of importin α/β-heterodimer formation.

## Results

### R-DPRs bind importin β and inhibit nuclear import

Although importin β has been shown to co-precipitate with poly-GR and poly-PR, a direct interaction between R-DPRs and importin β has not been demonstrated, and the consequences for functional nuclear import are unknown. To test for an interaction between *C9orf72* DPRs and importin β, we used a variant of the FRET sensor Rango (“<u>Ran</u>-regulated importin β car<u>go</u>”) (Kalab et al., 2006), which consists of the importin β-binding domain (IBB) of importin α1 (KPNA2), flanked by CyPet (donor) and YPet (acceptor). When bound to importin β (KPNB1), Rango FRET is low, but in the presence of RanGTP, importin β is displaced from the sensor and FRET increases (Figure 1A-C). Since conserved arginine and lysine residues of the IBB domain are required for binding to importin β (Görlich et al., 1996; Weis et al., 1996; Cingolani et al., 1999), we hypothesized that the arginine-rich DPRs could bind to the corresponding sites on importin β and compete with the IBB. Synthetic GP10, GA10, and PA10 peptides did not affect Rango FRET even at high concentrations (Figure 1D). However, we observed a dose-dependent increase in FRET with low-nanomolar PR10 and GR10, indicating these DPRs are capable of binding to importin β and displacing the sensor. To further validate these observations, we used GFP nanobody-coated beads to bind Rango and probe for co-immunoprecipitation of importin β in the presence of increasing concentrations of GR10 and PR10 (Figure 1E-F). Again, we observed the dose-dependent displacement of importin β from the sensor at low nanomolar concentrations, confirming that Rango release was responsible for the increases in FRET.

**Figure 1.**
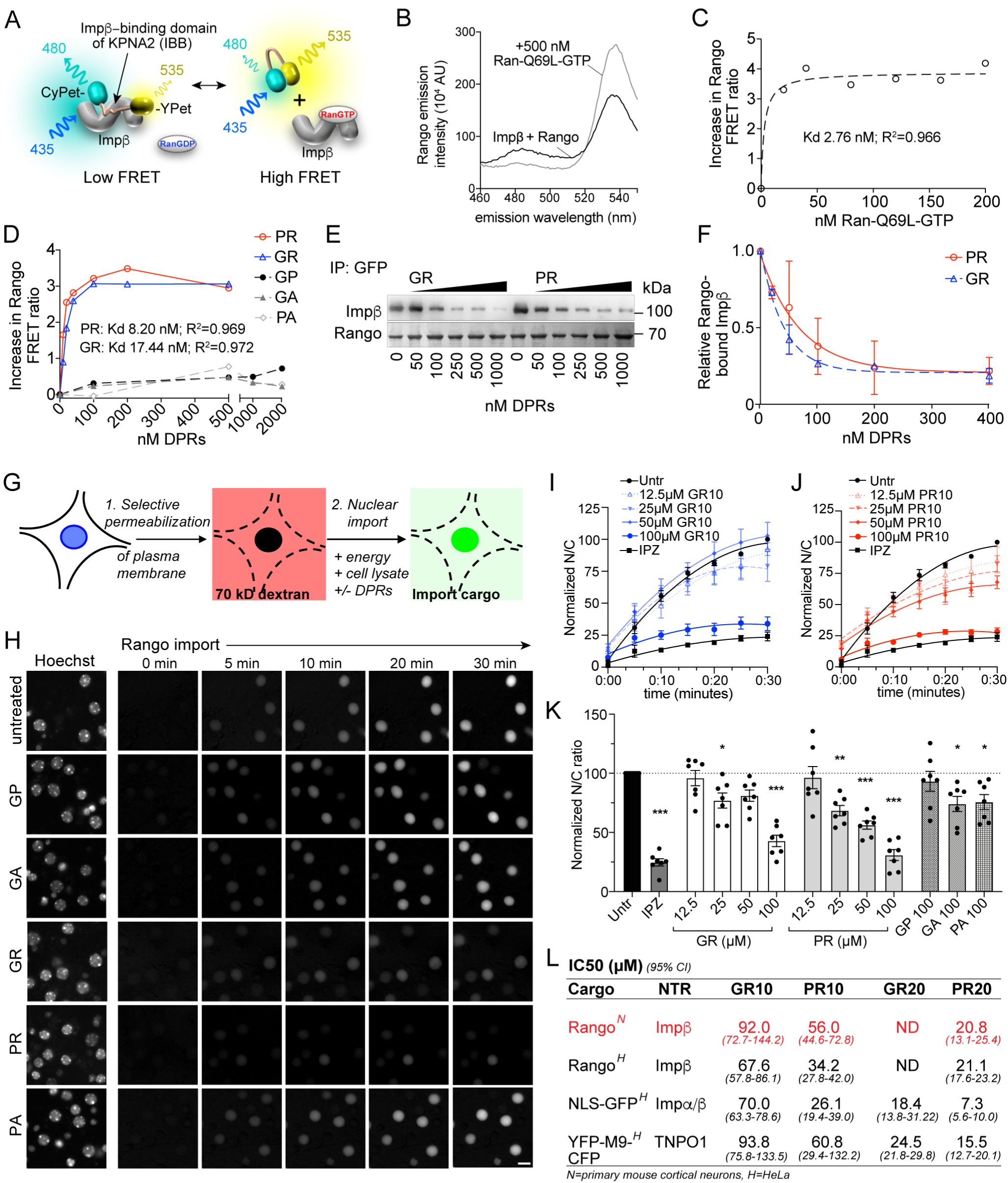
R-DPRs bind importin β and inhibit nuclear import. **A.** Schematic of Rango FRET sensor, consisting of the importin β-binding domain (IBB) of importin α1 (KPNA2), flanked by CyPET (donor) and YPet (acceptor). **B-C.** Rango spectral profile **(B)** and FRET ratio **(C)** demonstrating increase in FRET by adding hydrolysis-deficient Ran-Q69L-GTP to importin β-bound Rango (representative of 3 experiments). **D.** Change in Rango FRET ratio induced by adding DPRs (10-mers) to importin β-bound Rango (representative of 5 experiments, data in C-D fit to non-linear model with one binding site). **E.** GFP-trap co-immunoprecipitation of importin β by Rango in the presence of GR10 and PR10. **F.** Quantification of bound importin β in **(E)**, normalized to Rango and expressed as a fraction of untreated lysate (mean ± SD, two technical replicates). **G.** Diagram of permeabilized cell nuclear import assay, which was adapted and validated for primary neurons (**Figure 1-figure supplement 1**). **H.** Longitudinal wide-field images of Rango import in permeabilized mouse primary cortical neurons. Scale bar=10µm. **I-J.** Nuclear to cytoplasmic (N/C) ratio of Rango import in (**H**), calculated by automated high content analysis. GR and PR graphs are separated for clarity; the control values are identical. All data are normalized to cells lacking energy/lysate and expressed as percent untreated controls (mean ± SEM of n=4 biological replicates, 189 ± 125 cells per data point). **K.** Steady state N/C ratio of Rango in primary neurons fixed after 2 hours (mean ± SEM of n=7 biological replicates, 409 ± 202 cells per data point, \**p*<0.05, \*\**p*<0.01, \*\*\*\**p*<0.001 vs. untreated cells, one-way ANOVA with Dunnett’s post hoc test). **L.** IC50 of R-DPRs for inhibition of nuclear import of designated cargoes, from **(K)** and **Figure 1-figure supplement 2**. 95% confidence intervals are shown (n=3-6 biological replicates/condition, 409 ± 202 cells/ replicate for neurons, 1290 ± 305 cells/replicate for HeLa). See source file for raw data and exact *p* values.

To test the functional consequence of R-DPR-importin β interactions for nuclear import, we performed the permeabilized cell assay (Adam et al., 1990), in which the plasma membrane of cultured cells is selectively permeabilized, leaving the nuclear membrane intact as verified by nuclear exclusion of 70 kD dextran (Figure 1G). Fluorescent transport cargo is then added, with energy regeneration mix and cell lysate to provide a source of importins and Ran for nuclear import, which is measured by increasing nuclear fluorescence. Traditionally, this method uses digitonin for permeabilization; however, when attempted with primary mouse cortical neurons, we repeatedly found that even minimal concentrations of digitonin opened both the plasma and nuclear membranes. Since the nuclear envelope is devoid of the digitonin target cholesterol (Colbeau et al., 1971; Adam et al., 1990), we reasoned that its rupture in permeabilized neuronal cells was caused by mechanical perturbation upon removal of cytoplasmic proteins. Therefore, we developed a new protocol involving hypotonic cell opening in the presence of a high concentration of BSA as a cushion, which facilitated the selective plasma membrane opening of neurons (Figure 1-figure supplement 1).

Using this method, we performed live imaging of nuclear import of Rango, a direct importin β cargo whose Ran-, importin β-, and energy-dependent nuclear translocation is conferred by the IBB domain (Kalab et al., 2006). We verified that Rango import in permeabilized neurons is indeed dependent on energy and cell lysate, and can be inhibited by the importin β small molecule inhibitor, importazole (IPZ) in primary cortical neurons (Figure 1-figure supplement 1) (Soderholm et al., 2011). Time-lapse imaging of Rango import for 30 minutes in permeabilized neurons showed no effect of GP10, GA10, or PA10 at up to 100 µM, whereas GR10 and PR10 showed dose-dependent inhibition of transport (Figure 1H-J). The reaction was allowed to reach steady-state and fixed at 2 hours, at which point we observed statistically-significant transport inhibition beginning at 25 µM for both GR and PR (Figure 1K), with estimated IC50s as shown in figure 1L. In contrast, only trace inhibition by GA10 and PA10 was seen even at 100 µM, and there was no effect of 100 µM GP10. To facilitate testing of a broader range of cargoes and concentrations, we performed the assay in HeLa cells, with similar results to those seen in neurons (Figure 1L and Figure 1-figure supplement 2). To verify that the behavior of Rango in nuclear import signals indeed corresponds to endogenous importin α/β complexes, we tested the effect of DPRs on import of GST-GFP-NLS (hereafter referred to as GFP-NLS), a similarly-sized cargo that is loaded on importin β-bound importin α. Consistent with the expected lower efficiency of tripartite nuclear import complex assembly, R-DPRs perturbed GFP-NLS import even more strongly than that of Rango (Figure 1L and Figure 1-figure supplement 2).

The mechanisms of cargo recognition for importin β differ significantly even from its structurally closest relative TNPO1 (KPNB2), whose cargos are marked by the PY-NLS motif (Lee et al., 2006). However, since the sequence of the PY-NLS also contains basic residues, we tested the effect of R-DPRs on the nuclear import of YFP-M9-CFP (hereafter referred to as M9), a TNPO1 substrate based on the prototypic PY-NLS sequence of hnRNPA1 (Siomi and Dreyfuss, 1995) (Figure 1L and Figure 1-figure supplement 2). All substrates showed selective inhibition by GR and PR, which was more potent for PR and approximately 3-fold more potent on average for 20mers than 10mers. These results confirm that R-DPRs inhibit both importin β- and TNPO1-mediated nuclear import in this *in vitro* model system.

### R-DPRs interact with importin β in the bead halo assay

To further validate the direct interaction between R-DPRs and importin β, we performed the bead halo assay. This equilibrium-based binding assay is capable of identifying both low- and high-affinity interactions between ‘bait’ proteins immobilized to beads, and fluorescent ‘prey’ in the surrounding buffer, which forms a fluorescent halo on the bead surface (Patel et al., 2007; Patel and Rexach, 2008). First, we examined the propensity for all five DPRs to interact with biotinylated importin β, immobilized on the surface of neutravidin beads (Figure 2A). Controls included bare beads and beads coated with biotinylated BSA. As a positive control, we observed that the Rango sensor exclusively bound to full-length importin β-coated beads, and not the control beads. Fluorescent dextran, the negative control, did not form a halo in any conditions. AF488-labeled PR10 and GR10 (200 nM) both showed a modest degree of non-specific binding to all controls which was equivalent to the binding seen to bare beads. However, there was an approximately two-fold more intense halo around importin β-coated beads versus controls (Figure 2B), as quantified by the ratio of the fluorescent rim of the beads (the intensity around the surface of the beads at their equator) to the background fluorescence (Figure 2-figure supplement 1). When we added 1 mg/ml neuronal lysate to test the stringency of the interaction, all binding between GR10 and the beads, including importin β, was lost (Figure 2C-D). For PR10, nonspecific binding decreased, but the intensity of the importin β halo increased. These findings further support a direct interaction between R-DPRs and importin β, while indicating a higher relative selectivity of PR for importin β, compared to GR.

**Figure 2.**
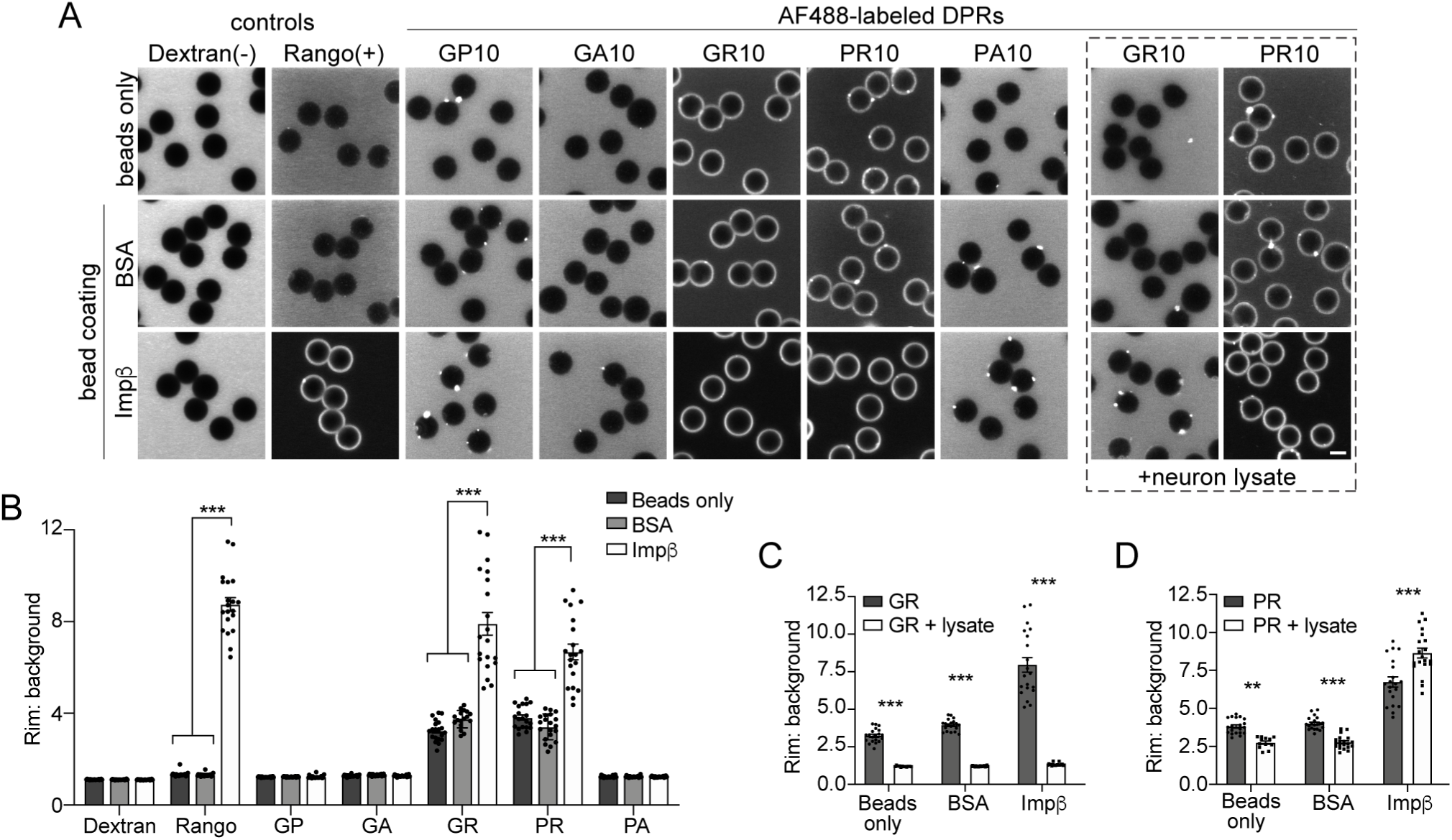
R-DPRs bind importin β in the bead halo assay. **A.** Confocal images of AF488-labeled *C9orf72* DPRs added to neutravidin beads coated with biotinylated ‘bait’ proteins, in binding buffer or in the presence of 1 mg/ml neuron lysate (at right). FITC-dextran = negative control (-), Rango sensor = positive control (+). Scale bar = 4µm**. B.** Rim vs. background ratio in binding buffer (see **Figure 2-figure supplement 1** for quantification method). **C-D.** Rim vs. background ratio for GR10 **(C)** and PR10 **(D)** in 1 mg/ml neuron lysate. In **B-D,** mean ± SEM is shown for n=20 beads (5 intensity profiles/bead). \*\**p*<0.01, \*\*\**p*<0.001 vs. control beads by two-way ANOVA with Tukey post-hoc test. See source file for raw data and exact *p* values.

### R-DPRs accelerate passive nuclear influx

To test if the disruption of nuclear import resulted from changes in the passive exclusion limit of NPCs, we tested the effects of R-DPRs on the passive influx of small cargoes. Passive diffusion of GFP and small fluorescent dextrans into nuclei of permeabilized HeLa cells was imaged at 10-second intervals for 5 minutes, and nuclear fluorescence quantified over time. All experiments were done in the context of energy and cell lysate, identical to the active transport conditions, so as not to miss putative effects that may depend on simultaneous active transport (i.e., recruitment of importins and DPRs to the NPC). Under these conditions, we observed the expected differences in the rates of passive influx of 10-, 40-, and 70-kD dextrans, and verified that addition of energy and lysate did not affect the baseline rate of passive influx of GFP (27 kD, no NLS) (Figure 3-supplemental figure 1). When we preincubated permeabilized nuclei with high concentrations of R-DPR 10mers or 20mers for 30-60 minutes, we observed no slowing of passive nuclear influx (Figure 3). Instead, R-DPRs accelerated the nuclear influx of both GFP and 40-kD dextran.

**Figure 3.**
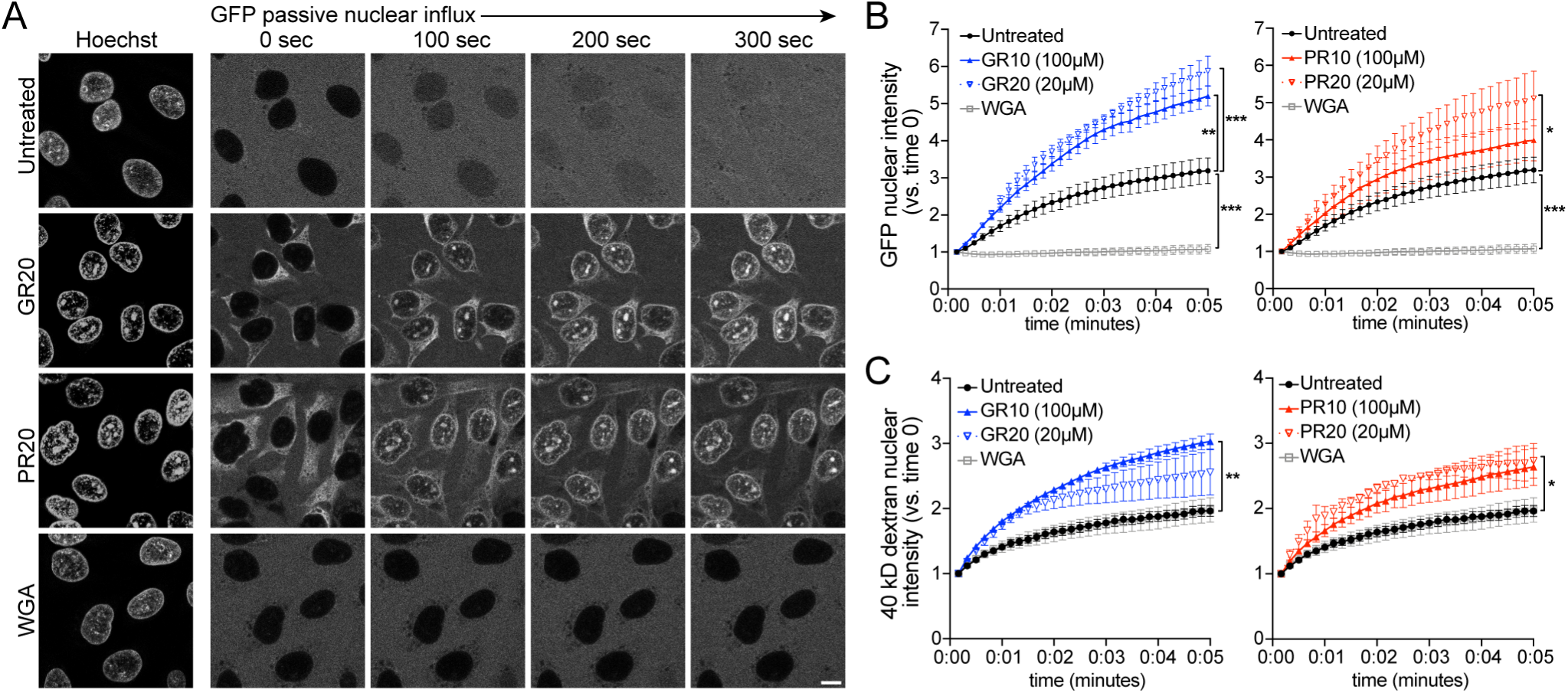
PR and GR accelerate passive nuclear influx. **A.** Confocal time-lapse imaging of GFP nuclear influx in permeabilized HeLa cells following ≥ 30 min. incubation with buffer (untreated), 20 µM GR20, 20 µM PR20, or 0.8 mg/ml wheat germ agglutinin (WGA, positive control). Scale bar = 10µm. **B-C.** Nuclear GFP (**B**) and 40 kD dextran (**C**) intensity normalized to background fluorescence, expressed vs. time 0 (no influx = 1). GR and PR are separated for clarity; the control values are identical. All experiments included lysate and energy, see **Figure 3-figure supplement 1** for validation of assay conditions, and **Figure 3-figure supplement 2** for binding studies with FG-domains which contribute to the NPC selectivity barrier. Data are mean ± SEM for n=3-6 biological replicates/condition (20-30 cells/replicate). \**p*<0.05, \*\**p*<0.01, \*\*\**p*<0.001 vs. untreated cells at 5 minutes by one-way ANOVA with Dunnett’s post hoc test. See source file for raw data and exact *p* values.

The rate of passive transport is thought to be governed by the FG-Nup barrier in conjunction with importin β and RanGTP (Ma et al., 2012; Kapinos et al., 2017). PR20 was previously shown by super-resolution microscopy to localize to the central channel of *Xenopus* oocyte NPCs, where it was hypothesized to inhibit both passive and active nuclear transport via stabilization of FG-domains (Shi et al., 2017). To verify DPR-FG binding, we used the bead halo assay to probe for interactions between all five *C9orf72* DPRs and yeast FG and GLFG domains (Figure 3-supplemental figure 2). As in the importin β halo assay, we observed moderate non-selective binding by the R-DPRs to all beads, including those coated with an F->A mutant construct. However, quantification of the halo intensities showed additional selective binding of both GR10 and PR10 to FG-domains of Nup100 (yeast homolog of Nup98), but not Nsp1 (yeast homolog of Nup62). For PR10, FG-binding could be augmented (to both Nup100 and Nsp1 fragments) by adding unlabeled importin β to the assay, suggesting that recruitment of PR to FG domains at the NPC could be mediated in part by an indirect interaction through importin β. Overall, these results support direct and indirect binding of R-DPRs to FG domains. Importantly, based on our passive transport studies, these interactions do not confer an impedence to transport as previously suggested, but rather a modest increase in NPC permeability.

### R-DPR-induced aggregates recruit NCT proteins

Upon addition of R-DPRs to cell lysate for the transport assays, we observed the rapid formation of insoluble aggregates (Figure 4A). To identify the components of these aggregates and determine their potential relevance for the nuclear import defect, we spun them down and analyzed their protein content via mass spectrometry (Figure 4A-B; data uploaded to http://proteomecentral.proteomexchange.org). 858 proteins were identified in each of two GR replicates and 758 in two PR replicates, with 647 (67%) in common. Consistent with previous reports, these included numerous nucleic acid-binding proteins and ribosomal subunits. Gene ontology (GO) analysis confirmed enrichment of nucleolar proteins, ribonucleoproteins, spliceosomal complex subunits, stress granule constituents, and others (Figure 4-figure supplement 1). Among these, low complexity domain (LCD)-containing proteins implicated in ALS/FTD were identified including TDP-43, FUS, Matrin-3, and hnRNPs. Multiple NCT proteins including karyopherins, Nups, Ran cycle proteins, and THO complex proteins, which participate in mRNA biogenesis and nuclear export (Rondón et al., 2010), were also found among the identified targets (Figure 4A-B).

**Figure 4.**
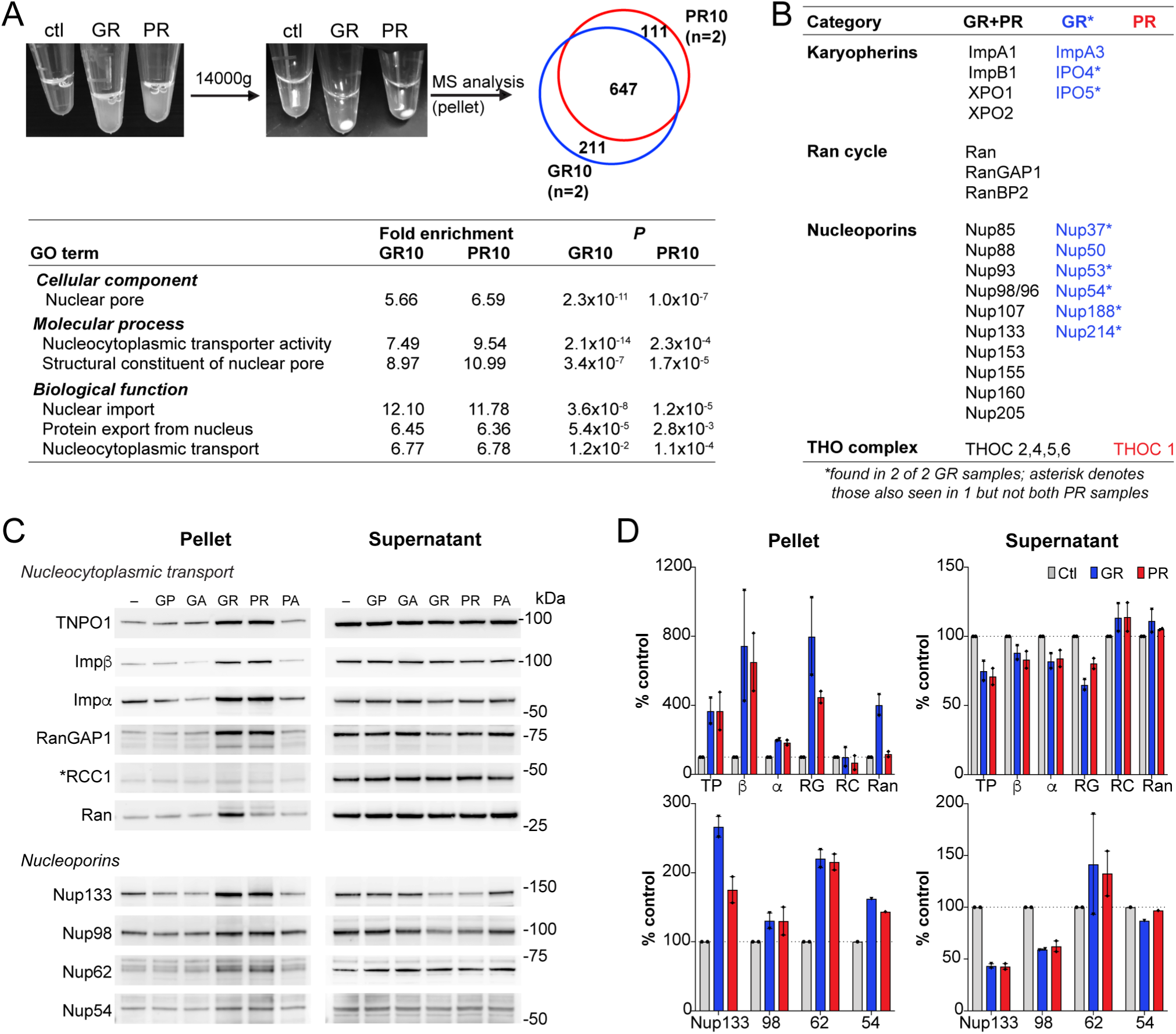
R-DPR-induced aggregates recruit NCT proteins. **A.** Aggregates formed by adding R-DPRs to HEK cell lysate in transport buffer (before and after 15 min centrifugation). Venn diagram indicates number of proteins identified by mass spectrometry analysis of pellets (n= 2 technical replicates). Enriched NCT-related GO terms are shown, with fold change and *p* value calculated by the DAVID algorithm. Overall top GO terms are shown in **Figure 4-figure supplement 1**. **B.** List of identified NCT-related proteins, in all 4 samples (black), n=2 GR10 samples (blue), and n=2 PR10 samples (red). Asterisk denotes samples seen in n=2 GR10 samples and only n=1 PR10 sample. **C.** Western blots for indicated NCT and Nup proteins in pellet vs. supernatant fractions. RCC1 is marked with an asterisk, as this protein was not identified in the MS results and serves as the negative control. All samples were loaded by volume, see **Figure 4-figure supplement 2** for membrane protein stain and additional Western blots of disordered RNA binding proteins**. D.** Quantification of blots in (**C**). Mean ± SD for two technical replicates is shown (TP=TNPO1, β=importin β, α=importin α, RG = RanGAP1, RC = RCC1, Ran = RanGTPase). See source file for raw data.

Next, we validated a subset of these identified proteins by Western blot, focusing on NCT proteins, Nups, and LCD-containing proteins (Figure 4C-D and Figure 4-figure supplement 2). We compared supernatant versus pellet fractions for all five DPRs compared to control lysates in which no DPRs were added, to assess the degree to which proteins were being sequestered and depleted from the soluble fraction. We saw enrichment in the pellet for importin β, RanGAP1, TNPO1, Ran, and importin α, with only minor decreases in the supernatant. RCC1 was not identified by mass spectrometry, and as predicted did not sediment with the DPRs, serving as a negative control. We also confirmed deposition of nucleoporins 54, 62, 98, and 133 in the pellet (Figure 4C-D), along with the low complexity domain (LCD)-containing RNA binding proteins TDP-43, FUS, Matrin-3, hnRNP A1, and hnRNP A2/B1, ribosomal protein RPS6, and the ATP-dependent RNA helicase DDX3X (Figure 4-figure supplement 2). As opposed to the NCT proteins, many of these LCD-containing proteins were markedly or completely depleted from the supernatant.

These data confirm that R-DPR aggregates can recruit NCT constituents in addition to a host of nucleic acid-binding proteins. However, NCT proteins were not substantially depleted from the supernatant even in the presence of 100 µM GR10 and PR10, suggesting that sequestration of critical NCT factors in these insoluble protein assemblies is unlikely to fully explain the failure of nuclear import in the transport assays.

### R-DPR nuclear import blockade does not require aggregates and is rescued by RNA

Cytoplasmic aggregate formation, a well-known pathological hallmark of neurodegenerative disease, has been proposed as a general mechanism for impairment of NCT (Woerner et al., 2016), although there is no evidence that such accumulation alters or disorganizes the NPC. It is unclear whether it is the disordered proteins themselves, or the process of aggregate formation, that may disrupt NCT. To address this question in the context of R-DPR aggregates, we tested several approaches for preventing aggregate formation in our model system. Addition of the aliphatic alcohol, 1,6-hexanediol, previously shown to disrupt GR- and PR-induced protein assemblies (Lee et al., 2016), was incompatible with transport and caused dose-dependent inhibition at baseline (Figure 5-figure supplement 1). This is likely due to disruption of FG-domains within the central channel, as previously reported (Ribbeck and Görlich, 2002). NTRs themselves, as hydrophobic interactors of aggregation-prone RNA binding proteins, have been shown to promote solubility of their cargoes and may have evolved in part as cytoplasmic chaperones (Jäkel et al., 2002; Guo et al., 2018; Hofweber et al., 2018; Yoshizawa et al., 2018; Qamar et al., 2018). However, even low concentrations of exogenous, full-length importin β inhibited nuclear import when added to the transport assay, likely due to sequestration of Rango and available RanGTP. Moreover, neither 1,6-hexanediol nor exogenous importin β could reverse mild nuclear import inhibition due to 25 µM PR10 (Figure 5-figure supplement 1).

Next, we tested the effect of increasing the concentration of RNA, based on the growing evidence that RNA is an important factor in mediating solubility of RNA-binding proteins (Maharana et al., 2018; Hondele et al., 2019; Mann et al., 2019). When total HEK cell RNA was added to the transport reaction, we saw a dose-dependent rescue of the import defect that was RNAse-sensitive (Figure 5A). However, this did not appear to be attributable to significant reduction of the quantity of insoluble material in the reaction (Figure 5-figure supplement 2). Instead, electrophoretic mobility shift assays showed that in a purified system, the RNA binds directly to the DPRs (Figure 5-figure supplement 2).

**Figure 5.**
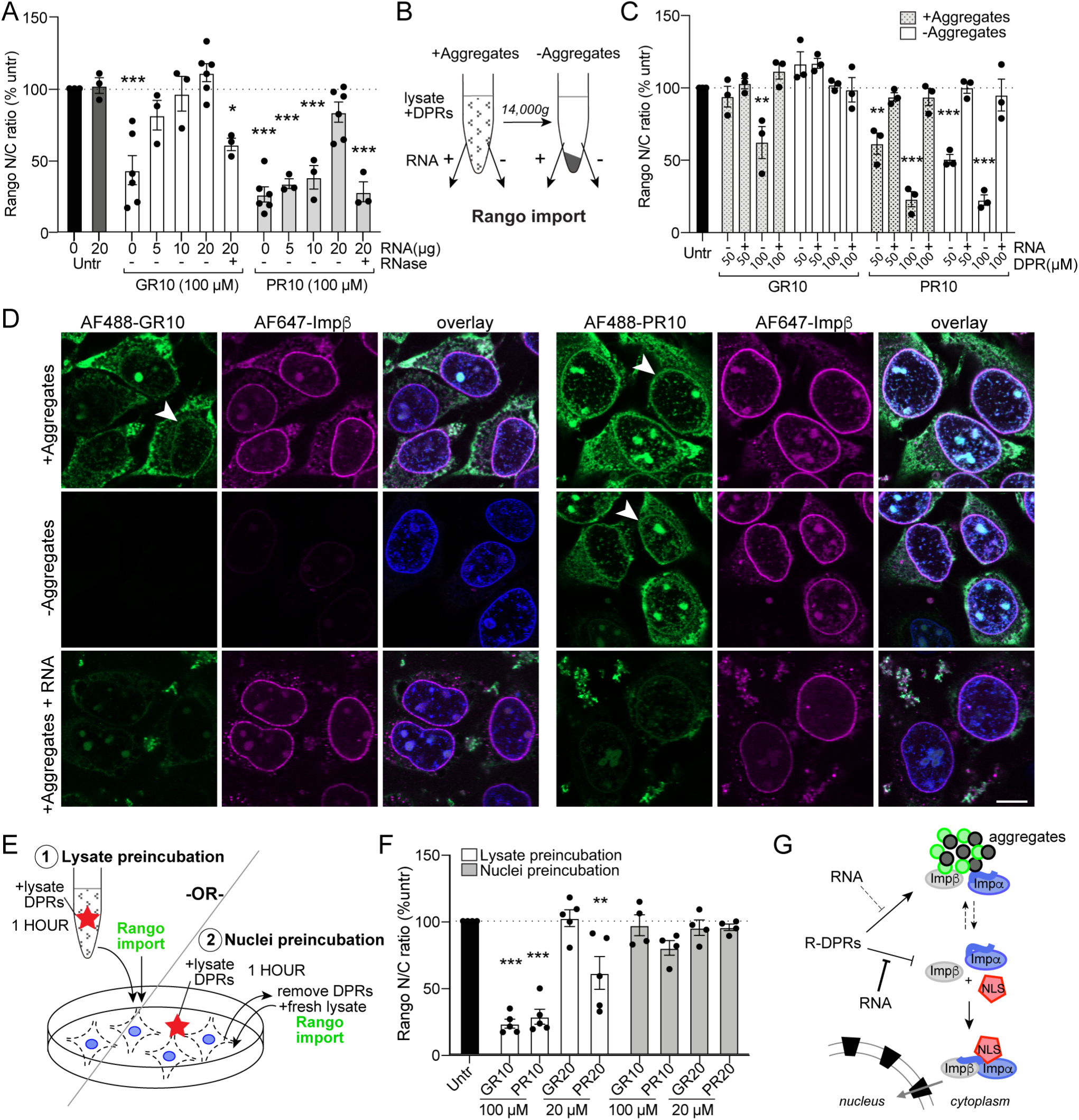
R-DPR nuclear import blockade does not require aggregates and is rescued by RNA. **A.** Rango N/C ratio in permeabilized HeLa transport reactions with 100 µM GR10 or PR10 and increasing concentrations of total HEK cell RNA +/− RNAse. **B.** Schematic of fractionated Rango transport assays, run with aggregates present or absent (supernatant only), followed by addition of RNA to a subset of reactions. **C.** Rango N/C ratio from fractionated transport assays. **D.** Confocal images of fractionated transport assays run in the presence of AF488-labeled R-DPRs and AF647-labeled importin β. Arrows mark R-DPR collection around the nuclear membrane in conditions where transport was inhibited. Acquisition parameters were kept constant for all images (scale bar=10 µm). **E.** Schematic of (1) lysate vs. (2) nuclei R-DPR preincubation assays. **F.** Rango N/C ratio from preincubation assays. **G.** Working model: R-DPRs block nuclear import by binding to importin β and preventing the formation of the importin α•importin β•NLS cargo complex in the soluble phase of the transport reaction, which can be alleviated by RNA. For **A,C,F** mean ± SEM of n≥3 biological replicates are shown (each data point represents 1462 ± 555 cells). \**p*<0.05, \*\**p*<0.01, \*\*\**p*<0.001 vs. untreated cells by one-way ANOVA with Dunnett’s post-hoc test. See source file for raw data and exact *p* values.

To further test if DPR-induced aggregate formation is relevant to the mechanism of nuclear import blockade, we performed a series of assays in which the supernatant was separated from the insoluble pellet prior to initiating transport (diagrammed in figure 5B). We reasoned that if aggregates sequester key transport factors, the remaining supernatant would be insufficient to drive nuclear import. However, if the aggregates contain inhibitor(s) of nuclear import or are themselves inhibitory, depleting them could rescue transport impairment. The results were markedly different for GR versus PR (Figure 5C). For GR10, removing the insoluble pellet restored nuclear import to normal, confirming that the inhibitory factor was present in (or was) the aggregates. In contrast, nuclear import remained perturbed in the supernatants of the PR10 aggregates, although it was restored by the addition of RNA.

Next, we monitored the location of the R-DPRs with respect to the aggregates by adding AF488-labeled DPRs to the transport reactions. By confocal microscopy, we observed that the transport disruption correlated with the presence of DPRs in the vicinity of the nuclear envelope (Figure 5D). AF488-GR10 fully sedimented into the pellet, leaving no visible GR10 in the supernatant, where transport proceeded normally. In contrast, a subset of AF488-PR10 remained in the supernatant and was present at the nuclear envelope, paralleling the persistent inhibition of nuclear import by the PR10 supernatants. RNA dispersed AF488-R-DPRs from the permeabilized cell nuclei in all conditions, restoring nuclear import. These results suggest that the import inhibition depends on GR or PR acting directly, rather than through putative intermediary factor(s), to inhibit nuclear import. The strikingly divergent segregation of GR10 vs. PR10 between the soluble and insoluble phase indicated that, while both share importin β as their target, the mechanisms and locations of their intracellular actions could differ significantly.

The critical steps of importin β-mediated nuclear import take place at NPCs via interactions with FG-Nups. To test whether interaction between R-DPRs and the NPC is sufficient to confer the block to import, we ran two parallel sets of import reactions (diagrammed in figure 5E). In the “lysate preincubation” paradigm, as for previous active import assays, R-DPRs were added to lysates used to supply transport factors, preincubated for 1 hour, and then added to permeabilized cells along with Rango and energy to initiate the transport reaction. In the “nuclei preincubation” set, we first exposed the permeabilized cell nuclei to R-DPRs (in the presence of lysate and energy, but no fluorescent cargo). After 1 hour, the DPR-lysate mix was removed from the nuclei, and fresh transport lysate, energy, and cargo added to initiate transport (without R-DPRs). We hypothesized that, if the DPRs inhibited Rango import by associating with and perturbing the NPC, we should see reduced import rate in the “nuclei preincubation” group. However, transport proceeded normally (Figure 5F). These results support a model in which the R-DPRs inhibit nuclear import by directly interfering with factor(s) present in the soluble phase of the NCT machinery (Figure 5G), which is consistent with the biochemical evidence for importin β as one of their direct targets.

## Discussion

Importin β, together with its importin α family of cargo adaptors, is fundamentally required for the nuclear import of NLS-containing proteins, including TDP-43, whose cytoplasmic mislocalization is observed in ≥97% of ALS cases, including *C9orf72*-ALS (Neumann et al., 2006; Mackenzie et al., 2014). Here, we demonstrate that *C9orf72* R-DPRs interact with importin β, which disrupts import complex formation and inhibits nuclear import in permeabilized cell assays. R-DPRs induce aggregation in the transport assay, including NCT proteins, although the association with aggregates does not substantially reduce the availability of critical components required for nuclear import. Rather, the transport blockade appears to depend on the ability of R-DPRs to interact with soluble nuclear transport receptors in the vicinity of the NPC, an effect which can be rescued by RNA, and is consistent with disruption of nuclear import complexes.

Members of the β karyopherin family have been consistently identified as genetic modifiers of *C9orf72* toxicity in fly, yeast, and human cell screens (Zhang et al., 2015; Freibaum et al., 2015; Jovičić et al., 2015; Boeynaems et al., 2016; Kramer et al., 2018), and shown to coprecipitate with R-DPRs (Lee et al., 2016; Lin et al., 2016; Yin et al., 2017). Using purified proteins, we demonstrate by FRET, bead halo, and co-immunoprecipitation, that R-DPRs bind importin β with low-nanomolar affinities. Micromolar concentrations were required to observe functional import blockade in the permeabilized cell assay, however 20mers were on average 3.3-fold more potent than 10mers across all cargoes, suggesting that longer DPRs, as are likely observed in patients, may be significantly more potent. The true length of R-DPRs in patients is unknown, although high molecular weight smears have been observed by SDS-PAGE (Zu et al., 2013). GGGGCC repeat lengths in the 1000s have been reported in postmortem brain (van Blitterswijk et al., 2013; Dols-Icardo et al., 2014; Nordin et al., 2015), although the processivity of ribosomes along the repeat RNA, and what terminates non-AUG translation, is unclear. The intracellular concentration of R-DPRs is also unknown. By ELISA, the poly-GP concentration in postmortem motor cortex has been estimated at a median of 322 ng/mg protein (Gendron et al., 2015), but comparable measurements for R-DPRs have been technically prohibitive in our hands to date.

Importin β is composed of 19 tandem HEAT repeats, coiled into a superhelix with exposed N-terminal RanGTP-binding domain and C-terminal importin α-binding domain (Cingolani et al., 1999). We predicted that R-DPRs might mimic the arginine- and lysine-rich IBB domain of importin α, forming an electrostatic interaction with acidic residues in the C-terminal domain of importin β. Indeed, R-DPRs displaced importin β from the importin α-IBB (Rango) FRET sensor at low nanomolar concentrations, supporting this hypothesis. At present, we cannot exclude binding to importin β at other sites, and given the similarity in HEAT repeat structure throughout the protein, multiple DPR-importin β interaction sites would not be surprising. Based on the propensity of R-DPRs to bind and induce aggregation of intrinsically-disordered proteins, the IBB domains of importins α and snurportin 1, both highly disordered unless bound to importin β (http://mobidb.bio.unipd.it/; Lott and Cingolani, 2011; Piovesan et al., 2018), could also be a target. It is conceivable that R-DPRs could perturb nuclear import by targeting both the IBB-binding site on importin β, and the IBBs on the much less abundant importins α (Görlich et al., 2003). Additional studies are needed to verify the precise domain(s) on importin β to which the R-DPRs bind, and to investigate direct importin α-targeting.

PR20 was previously shown to localize to the central channel of the NPC in Xenopus oocytes and inhibit nuclear import of NLS-BSA in permeabilized HeLa cells (Shi et al., 2017). In this study, the import blockade was attributed to binding and stabilization of FG domains in a polymerized state, creating a barrier to transport. We also observed modest R-DPR binding to FG-domains by the bead halo assay, as may be predicted by the ability of arginine-rich domains to undergo cation-pi interactions with aromatic phenylalanine rings. However, when we tested the functional consequences of R-DPRs on passive nuclear influx, a rate that depends both on the FG barrier and resident importin-cargo complexes (Kapinos et al., 2017), we observed marked acceleration. The precise cause is unclear, although the width of the passive channel has been shown to be modulated by the local concentration of importin β and RanGTP, increasing when importin β concentrations are high (Ma et al., 2012). Alternatively, leakiness of the passive barrier was reported in permeabilized cells where importin β/cargo complexes were depleted from the NPC (Kapinos et al., 2017). Additional studies are needed to test these possibilities. Nevertheless, incubation of permeabilized nuclei with high concentrations of R-DPRs blocked neither passive nor active transport. Taken together, these data do not support the hypothesis that alterations in the FG barrier account for R-DPR-mediated nuclear transport blockade.

Recent interactome screens have shown that R-DPRs engage in a broad range of protein-protein interactions (Lee et al., 2016; Lin et al., 2016; Yin et al., 2017) and can trigger phase separation of a large set of proteins involved in RNA metabolism and stress granule formation (Boeynaems et al., 2017). Even with 10mers, we similarly observed rapid alterations in the solubility of proteins upon adding R-DPRs to our transport reactions, that sedimented up to 10% of the cellular proteome and was markedly enriched for disordered nucleic acid-binding proteins. Aromatic rings common to many LCD-containing proteins, including FG-Nups, mediate their phase separation properties through cation-pi interactions with arginine residues (reviewed by Banani et al., 2017), and are likely perturbed by the rapid influx of R-DPRs. Indeed, *C9orf72* R-DPRs have been reported to disrupt phase separation properties of membrane-less organelles (Lee et al., 2016). While our MS analysis did not permit quantitative comparison between PR and GR, by Western blot we did observe varying selectivity for target proteins. Of note, a recent comparison between modifiers of R-DPR toxicity in yeast noted unexpectedly low overlap between GR- and PR-modifiers (Chai and Gitler, 2018), supporting the idea that they may behave differently in a physiologic context.

Based on growing evidence that RNA is integral to the solubility of disordered protein assemblies (Maharana et al., 2018; Langdon et al., 2018; Hondele et al., 2019), and polyU RNA can specifically coordinate the liquid liquid phase separation of PR (Boeynaems et al., 2017), we tested the effect of adding total cellular RNA to the transport reaction, and observed dose-dependent rescue. Total protein aggregates were not strongly reduced by the RNA, however significantly less AF488-labeled R-DPRs were observed in the vicinity of the nuclear envelope. Our electrophoretic mobility shift assay shows that a broad range of cellular RNAs can bind to R-DPRs directly, and previous evidence in a purified system showed that synthetic RNAs can facilitate suspension of R-DPRs in a droplet-like state (Boeynaems et al., 2017). The direct sequestration of R-DPRs by RNA could contribute to the reduced deposition of AF488 DPRs along the nuclear envelope, and the beneficial effects on nuclear import. At the same time, RNA could act indirectly to sequester the R-DPRs away from importins, by promoting the solubility of abundant disordered and RNA-binding proteins, thus decreasing aggregate size and increasing the availability of other R-DPR binding sites. While future studies will be needed to fully elucidate the mechanisms of direct and indirect effects of RNA on R-DPRs, our data suggest that, at least in the permeabilized cell model, RNA can mitigate aberrant protein-protein interactions in a functionally meaningful way.

In summary, we propose a model in which R-DPRs bind and interfere with nuclear import complex formation in the soluble phase of the nuclear transport reaction. Based on these findings, we speculate that importin β disruption may contribute to pathological protein mislocalization in *C9orf72*-mediated ALS/FTD, including TDP-43, for which links to downstream neurodegeneration are beginning to be unraveled (Ling et al., 2015; Melamed et al., 2019; Klim et al., 2019). Further investigation is warranted regarding disruption of importin β and other karyopherins in *C9orf72* disease models, and the potential for use of RNA-based strategies to mitigate aberrant R-DPR protein-protein interactions.

## Materials and methods

### DPR synthesis

10- and 20-mer dipeptide repeat proteins with C-terminal lysine (for solubility) and cysteine (for fluorescent tagging, i.e. GPGPGPGPGPGPGPGPGPGPKC) were synthesized by Genscript (Nanjing, China) and 21^st^ Century Biochemicals (Marlborough, MA) and verified by mass spectrometry to be free of trifluoroacetic acid adducts. Lyophilized powder was diluted in 0.1x XB’ buffer (5mM sucrose, 10mM KCl, 1 mM HEPES, pH 7.7) and frozen in single use 10 mM aliquots at −80°C after snap freezing in liquid nitrogen.

### Cloning of recombinant constructs

Restriction cloning was used to insert the ORF from pQE-ZZ-RanQ69L (Nachury and Weis, 1999) between the BamH1 and HindIII sites in pRSET A, resulting in pRSET ZZ-RanQ69L. The pRSET zzRCC1 was created by inserting the PCR-amplified wild-type (WT) human RCC1 C-terminally of the ZZ-tag in pRSET A. Site-directed mutagenesis and PCR cloning were used to modify Rango-2 (Kalab and Soderholm, 2010) by removing the KPN1 sites from YPet and CyPet (Nguyen and Daugherty, 2005) and replacing the Snurportin-1 IBB with the IBB amplified from human importin α1 (KPNA2). While doing so, the IBB-importin α1 domain was inserted either with (pK44) or without (pK188) flexible GGCGG linkers added between the 5’ and 3’ ends of IBB and the fluorophores. Restriction cloning was used to combine the C-terminal biotin acceptor peptide tag Avitag (GLNDIFEAQKIEWHE) from pAC-6 (Avidity, Aurora, CO) with WT human importin β (Chi et al., 1997) in pRSET A vector, resulting in in pRSET importin β-Avitag (pKW1982; pK1099). Restriction cloning in the modified pRSET A with C-terminal Avitag was used to create pRSET-EGFP-Avitag (pK803). The pGEX-2TK1 plasmid for the expression of the *S. cerevisiae* Nsp1(497-608) FxFG domain was obtained from M. Rexach (Yamada et al., 2010).

**Table 1.** Recombinant DNA constructs.

| DNA construct name | Reference or source |
| --- | --- |
| pRSET zzRanQ69L (pKW1234; pK1097) | this paper |
| pRSET zzRCC1 (pKW1907; pK1098) | this paper |
| pRSET Rango-2/α1+linkers (pK44) | this paper |
| pRSET Rango-2/α1 (pK188) | this paper |
| pRSET GFP-Avitag (pK803) | this paper |
| pRSET Importin β- Avitag (pKW1982; pK1099) | this paper |
| pAC-biotin ligase | Avidity |
| pRSET YFP-M9-CFP (pKW1006) | Soderholm et al., 2011 |
| pET30a 6His-S-Importin $\beta$<sub>(1-876)</sub> (pKW485) | Chi et al., 1997 |
| pGEX GST-GFP-NLS (pMD49) | Levy and Heald, 2010 |
| pGEX-2TK-Nup100<sub>(1-610)</sub> | Onischenko et al., 2017 |
| pGEX-2TK-Nup100<sub>(1-307)</sub> | Onischenko et al., 2017 |
| pGEX-2TK-Nup116<sub>(348-458)</sub> | Onischenko et al., 2017 |
| pGEX-2TK-Nup116<sub>(348-458)</sub> F>A | Onischenko et al., 2017 |
| pGEX-2TK Nsp1<sub>(497-609)</sub> (pKW1609; pK1100) | Yamada et al., 2010 |

### Recombinant protein expression

Unless otherwise specified, recombinant proteins were expressed in *E. coli* BL21(DE3) cells (ThermoFisher, Waltham, MA) that were cultured in 1L batches of LB media contained in 2.8L baffle-free Fehrnbach flasks. Protein expression was induced with 0.3mM IPTG. Centrifugation was used to collect the cells and wash them in the ice-cold protein-specific buffer, as indicated below. Unless otherwise specified, all buffers were pH 7.4. The washed cell pellets were flash-frozen in liquid nitrogen and stored at −80°C, and lysed in ice-cold conditions and in the presence of protease inhibitors, using French pressure cell or microfluidizers. After dialysis in the protein-specific buffer, protein concentration was measured with the Bradford assay (BioRad, Hercules, CA), and single-use aliquots of all proteins were stored at −80°C after flash-freezing in liquid nitrogen.

#### Recombinant proteins with GFP-derived tags

For expression of proteins containing GFP variants, including Rango (pK44 and pK188), YFP-M9-CFP, GST-GFP-NLS, and GFP-Avitag, the cells were first outgrown at 37°C until reaching OD_600nm_=0.1-0.3. The cultures were cooled to room temperature (22-25°C), and protein expression was induced at 22-25°C for 12-14 h.

Cells expressing 6His-tagged fluorescent proteins (Rango pK44 and pK188, YFP-M9-CFP, and GFP-Avitag) were washed and lysed in 10mM imidazole/PBS and purified with either Ni-NTA agarose (Qiagen, Venlo, Netherlands) or HIS-Select HF Nickel Affinity Gel (Millipore Sigma, St. Louis, MO). The lysates were clarified (40 min, 16000g, 4°C) and incubated with Ni resin (30-60 min, 4°C). The resin was placed into small chromatography columns, washed with ice-cold 10mM imidazole/PBS, and the proteins eluted with increasing concentration of imidazole/PBS (25-300mM). SDS-PAGE was used to select and pool batches with the highest purity, prior to dialysis in PBS or XB buffer (50mM sucrose, 100mM KCl, 10mM HEPES, 0.1 mM CaCl_2_, 1mM MgCl_2,_ pH 7.7).

Cells expressing GST-GFP-NLS were washed and lysed with TBSE (50mM Tris, 150mM NaCl, 4mM EDTA, pH8.0), the lysate clarified, and the protein affinity-purified on glutathione sepharose (Roche, Basel, Switzerland). After washes with TBSE, the proteins were eluted with TBSE containing increasing concentrations of glutathione (2.5-10 mM). Proteins eluted with 2.5 and 5mM glutathione were pooled and dialyzed in PBS before storage.

#### FRET assay mix with importin β and Rango

Full length human importin β was expressed from pET30a-WT Importin β (pKW485; (Chi et al., 1997)) at the Protein Expression Laboratory (PEL, National Cancer Institute, Frederick, Maryland). The transformed BL21DE3 cells were grown at 37°C in an 80L Bioflow 500 bioreactor (New Brunswick Scientific, Edison, NJ) until OD600_nm_ = 0.6, cooled to 22°C and the expression was induced with 0.3mM IPTG. After 12h induction, cells were harvested with the CARR continuous flow centrifuge and lysed in PBS with 10mM imidazole and 5mM TCEP with a 110EH Microfluidizer (Microfluidics, Westwood, MA) using 2 passes at 10,000 PSI under chilled conditions. The lysates were flash-frozen in liquid nitrogen and stored at −80°C. After thawing, the lysates were clarified and incubated with HIS-Select HF Nickel Affinity beads. The beads were washed with 10mM imidazole/PBS and the protein eluted with 200mM imidazole/PBS before dialysis in XB. The purified importin β was combined with a freshly thawed aliquot of Rango2-α1 (pK188) at 2.5:1 molar ratio ratio (12.5 µM importin β, 5µM Rango) and supplemented with 3% glycerol. Measurement of Rango fluorescence emission in a spectrometer (see below) was used to verify the FRET-off state of the Rango/importin β mixture before freezing.

#### FRET assay mix with zz-RCC1 and zz-RanQ69L

The expression of zz-RCC1 was induced at OD_600nm_= 0.4, followed by incubation at 22°C for 4 hours. The cells were washed with PBS, 10mM Imidazole, 1mM MgCl_2_, 5mM TCEP, 0.2mM AEBSF, pH 8.0, and lysed by ice-cold microfluidizer. The clarified lysates were used to isolate the zz-RCC1 proteins on HIS-Select HF Nickel Affinity beads, as described above. Proteins eluted with 0.2M imidazole/PBS were dialyzed in PBS before storage. The zz-RanQ69L (pKW1234; pK1097) was expressed from BL21DE3 cells at PEL in an 80L bioreactor, using conditions described for importin β above, except that expression was induced with 0.3mM IPTG at 37°C for 3 hours, and lysis was performed in PBS with 10mM Imidazole, 5mM TCEP, 2mM MgCl_2_, and 1 mM GTP. The lysates were clarified and bound to HIS-Select HF Nickel Affinity beads (Millipore Sigma). The Ni resin was washed with ice-cold 10mM imidazole/PBS and the protein eluted with 0.2M imidazole/PBS, followed by dialysis in XB. After measuring the concentration, 60 µM zzRCC1 and 2.4 µM zzRCC1 were combined in XB containing 2mM GTP. Before aliquoting and storage, the measurement of Rango fluorescence emission in a spectrometer (see below) was used to verify that the zzRanQ69L-GTP-containing mix robustly induced Rango dissociation from importin β.

#### Importin β biotinylation

To prepare biotinylated WT importin β-Avitag (pKW1982, pK1099), BL21DE3 cells (New England Biolabs, Ipswich, MA) were co-transfected with the respective plasmids together with pAC-biotin ligase (Avidity), followed by plating and growth in LB media containing ampicillin and chloramphenicol. After the 37°C cultures reached OD_600nm_ + 0.4-0.6, the cultures were cooled to room temperature, supplemented with 100µM D-biotin, and the expression was induced with 0.3mM IPTG at room temperature for 8-11 hours (pKW762). Proteins were purified on Ni-NTA resin as described for the non-biotinylated importin β fragments.

#### FG- and GLFG-nucleoporin fragments

The expression of GST-Nsp1_(497-609)_ in BL21(DE3) cells grown in LB media was induced at OD_600nm_= 0.4-0.6, followed by incubation at 37°C for 3-5 hours. The GST-tagged *S. cerevisiae* pGEX-Nup100_(1-307),_ Nup100_(1-610)_, Nup116_(348-458)_ and Nup116_(348-458)_ F>A were expressed in T7 Shuffle cells (NEB) that were grown in Dynamite media (Taylor et al., 2017) until OD_600nm_= 0.9 before induction with IPTG at 37°C for 3 hours. All the GST-tagged Nup fragments were purified using glutathione-sepharose affinity chromatography, as described for the GST-GFP-NLS above, and dialyzed into PBS before storage.

### Importin β labeling with Alexa-647 and BSA biotinylation

Purified WT importin β (pKW485) diluted to 10 µM in XB was combined with 10-molar excess of Alexa Fluor 647 NHS ester (ThermoFisher), using freshly-prepared 10mM dye in anhydrous DMSO. After incubation on ice for 2 hours, the sample was dialyzed in PBS and concentrated on 30kD MWCO filter (Millipore Sigma) before storage. A similar protocol was used to label BSA with 10-molar excess Sulfo-NHS-LC-biotin (ThermoFisher), followed by dialysis in PBS.

### DPR labeling with Alexa-488-maleimide

Just before labeling, Alexa Fluor 488 C5-maleimide (ThermoFisher) was diluted to 20 mM in anhydrous DMSO and further diluted to 1.6 mM in XB’ buffer. Freshly thawed 10mM aliquots of DPRs were diluted to 2mM with 0.1x XB’ and combined with an equal volume of 1.6 mM Alexa Fluor 488 C5-maleimide and kept overnight at 4°C. The unreacted maleimide was quenched by 1:50 (v/v) 100mM DTT before aliquoting and storage at −80°C. To verify labeling, DPRs were separated by SDS-PAGE followed by fluorescence detection.

### Rango FRET detection

FRET assays for DPR-induced dissociation of the importin β-Rango complex were performed using a mix of 5 µM Rango-2/α1 (pK188) and 12.5 µM importin β (pKW485) prepared as described above. After thawing on ice, the mix was diluted to 20 nM Rango and 50 nM importin β in TBS, pH 7.4, 0.005% Tween-20 (TTBS), supplemented with increasing DPR concentrations, and mixed by brief vortexing at low speed. The positive control reactions for RanGTP-induced Rango-importin β dissociation were prepared by adding increasing concentrations of ZZ-RanQ69L-GTP to the samples, using a freshly thawed aliquot of 60 µM ZZ-RanQ69L, 2.4µM ZZ-RCC1, 2mM GTP in XB. The assay buffer alone was used as a blank. A Fluoromax-2 spectrometer (Jobin Yvon Horiba, Piscataway, NJ) was used to detect the Rango emission spectra (460-550nm, in 1nm increments) while exciting the samples at 435nm. The excitation and emission bandpass were set to 5nm and integration time to 0.05s. Peak emissions were recorded at 480nm (donor) and 535nm (acceptor) in all samples, and background emission subtracted at the same wavelengths in the blank. The FRET signal was calculated as the ratio of background-subtracted acceptor/donor emissions. The signal detected in the untreated sample (20 nM Rango and 50 nM importin β-only, the lowest FRET), was then subtracted from the resulting values. Prism v6 (Graphpad, San Diego, CA) was used to calculate the non-linear fit with one site-specific binding model while using the D’Agostino and Pearson K2 test to verify the normality of residuals and the Runs test to assure non-significant deviation from the model.

### Biochemical pulldown assay for DPR-induced Rango-importin β dissociation

An aliquot of 5 µM Rango-2/α1 + 12.5 µM importin β mix was diluted to 20 nM Rango and 50 nM importin β in TTBS, supplemented with increasing concentrations of GR10 or PR10, mixed by vortexing, and incubated for 30 min at room temperature. GFP-Trap magnetic beads (Chromotek, Planegg-Martinsried, Germany) were washed and resuspended in TTBS. At the end of incubation, 8µl bead suspension was added to each sample and mixed by rotation for 15 min. The supernatant was removed and beads washed 3 times with TTBS before boiling in 20µl SDS-PAGE sample buffer with 2% β-mercaptoethanol. Samples were separated by SDS-PAGE and anti-GFP Western blot performed as detailed below to detect Rango. After detecting the ECL signal, membranes were stained with Coomassie Brilliant R250 to detect importin β. Background-subtracted signals were determined by Image Lab 6.01 (BioRad) and the Rango ECL signal normalized to the importin β signal within each lane.

### Electrophoretic mobility gel shift assay for RNA-DPR interaction

Aliquots of total HEK RNA (3 µg) were mixed with either 4 µl 50µM DPR-AF488 in 0.1x XB’ or with 1 µl 0.2% SYBR Gold nucleic acid stain (ThermoFisher) diluted in water. After 5 min incubation at room temperature, the samples were supplemented with Fast Digest loading buffer (ThermoFisher; no nucleic acid stain) and separated by electrophoresis on native 1% agarose gel in TBE, alongside with lanes containing HEK RNA (3 µg) or RNA ladder mixed with SYBR Gold. Immediately after electrophoresis, the gels were photographed with Bio-Rad ChemiDoc XRS+ using UV transillumination to simultaneously visualize the AF488-labeled R-DPRs and SYBR Gold-labeled RNA signals (where added).

### Bead halo assay

The bead halo assay was carried out as described with minor modifications (Patel and Rexach, 2008), using 6-8 µM polystyrene beads coated with neutravidin (for biotinylated proteins) or glutathione (for GST-fusion proteins) (Spherotech, Lake Forest, IL). Beads were coated overnight at 4°C at saturating concentrations per manufacturers’ instructions and rinsed 2x in binding buffer (20 mM HEPES [pH 7.4], 150 mM KOAc, 2 mM Mg(OAc)_2_, 1 mM DTT, 0.1% Tween-20). Immediately prior to the assay, fluorescent bait proteins and beads were combined with 4x EHBN (40 mM EDTA, 2% 1,6-hexanediol, 40 mg/ml BSA, 500 mM NaCl) to a total of 40 µL per well, in optical glass-bottom 96 well plates (Cellvis, Mountain View, CA). Reactions were allowed to equilibrate at room temperature for a minimum of 30 minutes prior to imaging at 100x on an LSM800 confocal microscope (Zeiss, Oberkochen, Germany). Intensity profiles comparing the maximum rim intensity to the background were plotted in ImageJ (NIH) by an investigator blinded to experimental conditions.

### Mouse primary cortical neuron culture and permeabilization

All animal procedures were approved by the Johns Hopkins Animal Care and Use Committee. Timed pregnant C57BL/6J females (Jackson Laboratory, Bar Harbor, ME) were sacrificed by cervical dislocation at E16, cortex dissociated, and cells plated at 50,000/well on poly-D-lysine/laminin-coated, optical glass-bottom 96-well plates. Growth medium consisted of Neurobasal supplemented with B27, Glutamax, and penicillin/streptomycin (Gibco/ThermoFisher). At 5-7 days in vitro, neurons were rinsed in prewarmed PBS and permeabilized for 4 min. at 37° in a hypotonic solution containing 0-40 µM Tris-HCl pH 7.5 (to cause osmotic swelling) and 50-150 mg/ml BSA (for molecular crowding/mechanical support). Following permeabilization, cells were placed on ice and rinsed 2 x 5 minutes in transport buffer (TRB, 20mM HEPES, 110mM KOAc, 2mM Mg(OAc)_2_, 5mM NaOAc, 0.5mM EGTA, 250mM sucrose, pH 7.3, with protease inhibitor cocktail). All rinse and assay buffers were supplemented with 50 mg/mL BSA. The optimal hypotonic buffer and BSA concentration varied by batch, and was optimized prior to each set of assays for ability to permeabilize the majority of plasma membranes while maintaining nuclear exclusion of a 70 kD fluorescent dextran (ThermoFisher).

### HeLa cell culture and permeabilization

A single cell-derived clone of HeLa cells (ATCC, Manassas, VA; mycoplasma negative and validated by STR profiling) were maintained in OptiMEM (Gibco/ThermoFisher) with 4% FBS and plated on uncoated optical glass-bottom 96 well plates, at appropriate densities to reach 70-90% confluence on the day of the transport assay. To permeabilize, cells were rinsed for 2 minutes in ice-cold PBS, and permeabilized on ice for 10 minutes in 15-30 µg/mL digitonin (Calbiochem, San Diego, CA) in permeabilization buffer (PRB, 20mM HEPES, 110mM KOAc, 5mM Mg(OAc)2, 0.5mM EGTA, 250mM sucrose, pH 7.5, with protease inhibitor cocktail). Following permeabilization, cells were placed on ice and rinsed 3 x 5 minutes in transport buffer (TRB, 20mM HEPES, 110mM KOAc, 2mM Mg(OAc)_2_, 5mM NaOAc, 0.5mM EGTA, 250mM sucrose, pH 7.3, with protease inhibitor cocktail). The optimal digitonin concentration varied by cell density and passage number, and was optimized prior to each set of assays for the ability to permeabilize the majority of plasma membranes while maintaining nuclear exclusion of a 70 kD fluorescent dextran (ThermoFisher).

### Nuclear import assays

#### Assay components

Nuclear import was carried out essentially as described (Adam et al., 1990) with modified sucrose-containing buffers (Zhu et al., 2015). Concentrated whole cell lysates were prepared from HEK293T cells (ATCC, mycoplasma negative and validated by STR profiling), grown in 150 mm dishes and sonicated on ice in 1X TRB in the presence of protease inhibitor cocktail (Roche). The lysates were clarified (15 min, 14000g, 4C), snap frozen in liquid nitrogen, and stored in single use aliquots at −80C. Total HEK cell RNA was extracted using miRNEasy kits according to the manufacturers’ protocol, with DNase digestion (Qiagen). RNA concentration was measured by Nanodrop (ThermoFisher), and all 260/280 ratios were verified to be >2.0. Energy regeneration (ER) mix consisted of 100 µM ATP, 100 µM GTP, 4 mM creatine phosphate, and 20 U/mL creatine kinase (Roche).

#### Standard assay setup

Reaction mixes consisting of 2.5 mg/ml lysate, ER, fluorescent cargo (200 nM Rango and YFP-M9-CFP, 500 nM GST-GFP-NLS), Hoechst, DPRs, RNA, or inhibitors (100 µM importazole (IPZ, Millipore Sigma); 0.8 mg/mL wheat germ agglutinin (WGA, Millipore Sigma) were assembled on ice during cell permeabilization. DPRs or inhibitors were allowed to equilibrate in cell lysate for at least 30 minutes prior to initiation of transport. Every plate included: (1) Cargo alone: fluorescent cargo, but no ER or lysate, (2) Untreated controls: fluorescent cargo, ER, and lysate, and (3) Inhibitor: fluorescent cargo, ER, lysate, and IPZ (Rango and GST-GFP-NLS reactions) or WGA (YFP-M9-CFP). Preassembled transport reactions were then transferred onto permeabilized cells via multichannel pipette, and allowed to proceed at room temperature for 2 hours (Rango, YFP-M9-CFP) or 4 hours (GST-GFP-NLS). Cells were fixed in 4% paraformaldehyde/PBS, rinsed 2x with PBS, and transferred to 50% glycerol/PBS for immediate imaging.

#### Variations

For a subset of neuron transport assays, transport was monitored live via time lapse imaging every 5 minutes for 30 minutes. In a subset of HeLa assays, transport reactions were centrifuged before use at 14000g x 15 minutes to separate soluble and insoluble fractions. In another variation, the transport lysate + DPR and ER mix was allowed to preincubate on the permeabilized HeLa cells for at least 30 minutes prior to initiation of transport, rinsed 1x with TRB, and transport initiated with fresh lysate, cargo, and ER.

#### Imaging and data analysis

Multiple non-overlapping fields per well (4 for time-lapse imaging, 9-16 for fixed imaging), were captured at 40x on an ImageXpress Micro XLS high-content microscope (Molecular Devices, San Jose, CA), and the ratio of nuclear to cytoplasmic fluorescence intensity was calculated using the MetaXpress automated translocation-enhanced module. Raw data were filtered to exclude autofluorescence and the mean N/C ratio from wells without ER or cell lysate was subtracted from all values. Resulting N/C ratios were expressed as % untreated, to permit comparisons across biological replicates.

### Passive import assays

HeLa cells were permeabilized as above, rinsed 3 x TRB, and reaction mix containing 2.5 mg/ml HEK lysate, ER (to mimic the active import conditions), Hoechst, and/or DPRs were added and allowed to preincubate directly on the permeabilized cells for at least 30 minutes. 0.8 mg/ml WGA was used as a positive control. Cells were mounted on a Zeiss LSM800 confocal microscope, reaction mix withdrawn, and immediately replaced with fluorescent dextran (ThermoFisher) or recombinant GFP (pK803) in fresh lysate/ER mix to initiate the passive import reaction. A single 40x frame (containing 20-30 cells/well) was imaged per well, with images collected every 10 seconds for 5 min. The ratio of nuclear fluorescence intensity to local background at each timepoint was analyzed using Imaris (Bitplane, Zurich, Switzerland), and values for each cell were expressed as a ratio of time 0 (1 = no influx, >1 = influx).

### Mass spectrometry

50 µM GR10 or 25 µM PR10 (in duplicate) were added to 5 mg/ml HEK whole cell lysate (in TRB with ER), incubated for 60 min at 37°C and aggregates were pelleted by centrifugation at 16000g for 10 min. Supernatants were removed and pellets washed 2x and resuspended in MgCl_2_- and CaCl_2_-free DPBS (ThermoFisher), then flash-frozen in liquid nitrogen and stored at −80°C before further processing and analysis by the Johns Hopkins Mass Spectrometry and Proteomics core facility. Pellets were reduced/alkylated with DTT/IAA, reconstituted in TEAB/acetonitrile, and sonicated for 15 min prior to overnight digestion with Trypsin/LysC (Promega, Madison, WI) at 37°C. Some precipitate remained; supernatants were desalted and analyzed by LC/MS/MS on a QExactive_Plus mass spectrometer (ThermoFisher). MS/MS spectra were searched via Proteome Discoverer 2.2 (ThermoFisher) with Mascot 2.6.2 (Matrix Science, London, UK) against the RefSeq2017_83_ human species database (NCBI). Protein probabilities were assigned by the Protein Prophet algorithm (Nesvizhskii et al., 2003). Protein identifications were accepted if they contained at least 2 identified peptides at false discovery rate less than 1.0%. Gene ontology analysis was carried out using the DAVID algorithm v6.8 (May 2016, https://david.ncifcrf.gov/) (Huang et al., 2009a;b). The mass spectrometry data have been uploaded to the ProteomeXchange Consortium (http://proteomecentral.proteomexchange.org) via the PRIDE partner repository (Vizcaino et al., 2013), dataset identifier pending.

### DPR aggregation assay and Western blots

Supernatant and pellet fractions (for all 5 DPRs, and control/buffer only) were prepared as in the nuclear transport assays, by adding 100 µM 10mers to 2.5 mg/ml HEK lysate in 100 µL TRB. Supernatants were boiled in Laemmli (BioRad) for 5 minutes. Pellets were boiled for 15 minutes followed by sonication in order to fully disperse aggregates for SDS-PAGE. Equal volumes of supernatant and pellet fractions were run on 4-12% Bolt Bis-Tris Plus gels (ThermoFisher), transferred to nitrocellulose membrane using an iBlot2 dry blotting system (ThermoFisher). Protein loading was analyzed by BLOT-Faststain (G-biosciences, St. Louis, MO), according to the manufacterers’ instructions. For immunodetection, membranes were blocked with 5% non-fat milk in TBST and probed by sequential incubation with the primary antibodies as detailed in the table below. Detection was by HRP-conjugated secondary antibodies/chemiluminescence using an ImageQuant LAS 4000 system (GE, Chicago, IL). To permit sequential probing of membranes without stripping, signals were quenched by incubation with prewarmed 30% H_2_0_2_ for 20 minutes (Sennepin et al., 2009). Band intensities were measured by ImageQuant software. For pellet vs. supernatant fractions, all were expressed as percent untreated control. For blots in figure 1E, samples were run on 4-20% SDS PAGE minigels (ThermoFisher and blotted to PVDF membranes (Immun-Blot PVDF, Bio-Rad) using the Bio-Rad TransBlot Turbo apparatus, and probed as above. The chemiluminescence signal was captured with a Bio-Rad ChemiDoc XRS+ digital imaging system.

### Statistical analysis

Data analysis, graphing, and statistical analyses were carried out using Prism v6-v8 (Graphpad), according to methods detailed under each experimental approach above and in the figure legends.

### Antibodies

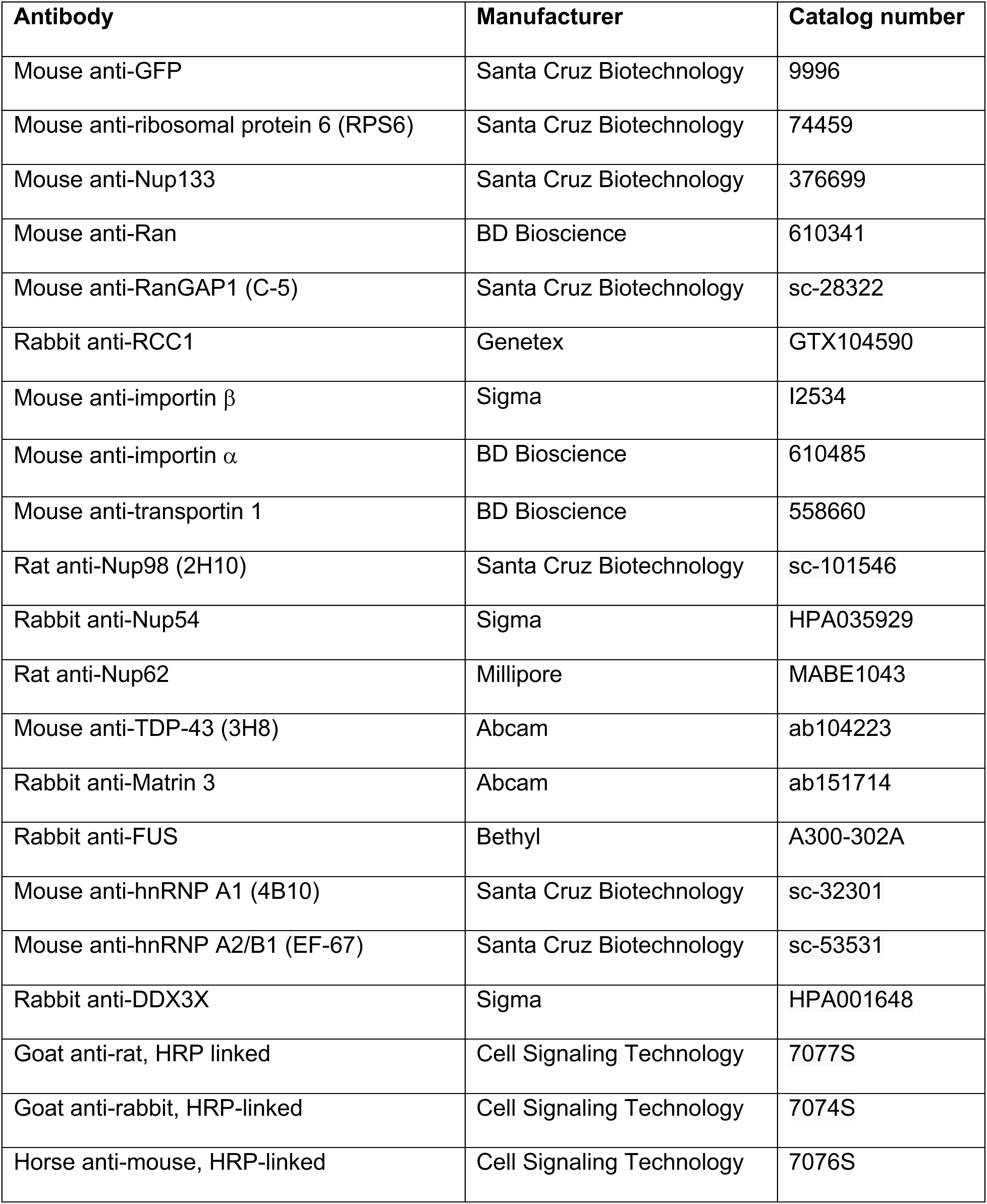

| Antibody | Manufacturer | Catalog number |
| --- | --- | --- |
| Mouse anti-GFP | Santa Cruz Biotechnology | 9996 |
| Mouse anti-ribosomal protein 6 (RPS6) | Santa Cruz Biotechnology | 74459 |
| Mouse anti-Nup133 | Santa Cruz Biotechnology | 376699 |
| Mouse anti-Ran | BD Bioscience | 610341 |
| Mouse anti-RanGAP1 (C-5) | Santa Cruz Biotechnology | sc-28322 |
| Rabbit anti-RCC1 | Genetex | GTX104590 |
| Mouse anti-importin $\beta$ | Sigma | I2534 |
| Mouse anti-importin $\alpha$ | BD Bioscience | 610485 |
| Mouse anti-transportin 1 | BD Bioscience | 558660 |
| Rat anti-Nup98 (2H10) | Santa Cruz Biotechnology | sc-101546 |
| Rabbit anti-Nup54 | Sigma | HPA035929 |
| Rat anti-Nup62 | Millipore | MABE1043 |
| Mouse anti-TDP-43 (3H8) | Abcam | ab104223 |
| Rabbit anti-Matrin 3 | Abcam | ab151714 |
| Rabbit anti-FUS | Bethyl | A300-302A |
| Mouse anti-hnRNP A1 (4B10) | Santa Cruz Biotechnology | sc-32301 |
| Mouse anti-hnRNP A2/B1 (EF-67) | Santa Cruz Biotechnology | sc-53531 |
| Rabbit anti-DDX3X | Sigma | HPA001648 |
| Goat anti-rat, HRP linked | Cell Signaling Technology | 7077S |
| Goat anti-rabbit, HRP-linked | Cell Signaling Technology | 7074S |
| Horse anti-mouse, HRP-linked | Cell Signaling Technology | 7076S |

### Reagents

| Reagent | Manufacturer | Catalog number |
| --- | --- | --- |
| GFP Trap Magnetic Agarose | Chromotek | Gtma-20 |
| Dynabeads MyOne Streptavidin C1 | ThermoFisher | 65001 |
| Sypro Ruby Protein Gel Stain | Millipore Sigma | S4942 |
| SYBR Gold Nucleic Acid Stain | ThermoFisher Scientific | S11494 |
| Importazole | Millipore Sigma | SML0341 |
| Alexa Fluor 647 NHS ester | ThermoFisher Scientific | A37573 |
| Alexa Fluor 488 C5 maleimide | ThermoFisher Scientific | A10254 |
| Biotin-maleimide | Millipore Sigma | B1267 |
| EZ-Link Sulfo-NHS-LC-biotin No-weigh format | ThermoFisher Scientific | A39257 |
| Ribo Ruler High Range RNA ladder | ThermoFisher Scientific | SM1821 |
| HIS-Select HF Nickel Affinity Gel | Millipore Sigma | HD537 |
| Ni-NTA Agarose | Qiagen | 30210 |
| Glutathione Sepharose 4B | GE Healthcare | 17-0756-01 |
| Zeba Microspin columns, 40 kDa MWCO, 75 µl | ThermoFisher Scientific | 87765 |
| BSA, fatty acid-free | Roche | 03117057001 |
| Glutathione-coated polystyrene particles 6.0-8.0 | SpheroTech | GSHP-60-5 |
| Neutravidin-coated polystyrene particles 6.0-8.0 | SpheroTech | NVP-60-5 |
| Dextran, Texas Red, 10,000MW | ThermoFisher Scientific | D1863 |
| Dextran, Texas Red, 40,000MW | ThermoFisher Scientific | D1829 |
| Dextran, Texas Red, 70,000MW | ThermoFisher Scientific | D1864 |
| Digitonin, high purity | Calbiochem | 300410 |
| Wheat germ agglutinin | Sigma | L0636 |
| miRNeasy kit | Qiagen | 217004 |
| DNase (RNase free) | Qiagen | 79254 |
| RNase A | ThermoFisher Scientific | EN0531 |

## Acknowledgements

Lin Xue and Svetlana Vidensky provided expert technical assistance. Robert Cole and Tatiana Boronina in the Johns Hopkins Proteomics Core assisted with proteomics analysis and interpretation. We gratefully acknowledge M. Rexach, M. Dasso, E. Onischenko and K. Weis for providing expression plasmids. This work was supported by NIH/NINDS (K08NS104273 [LH], R01NS094239 [JDR], P01NS099114 [JDR et al.],) and NIH/NIA (RF1AG062171 [JDR]). The funding sources had no role in study design, data collection and interpretation, or the decision to submit the work for publication.

## Author contributions

<u>LH</u>: conceptualization, methodology, validation, formal analysis, investigation, data curation, writing-original draft preparation, writing-review & editing, visualization, funding acquisition. <u>LD</u>: investigation, formal analysis, writing-review & editing. <u>KB</u>: investigation, formal analysis, writing-review & editing. <u>PK</u>: conceptualization, methodology, validation, formal analysis, investigation, resources, data curation, writing-review & editing, visualization, supervision. <u>JR</u>: conceptualization, writing-review & editing, supervision, project administration, funding acquisition.

**The authors declare no competing interests.**

---

**Figure 1-figure supplement 1.**
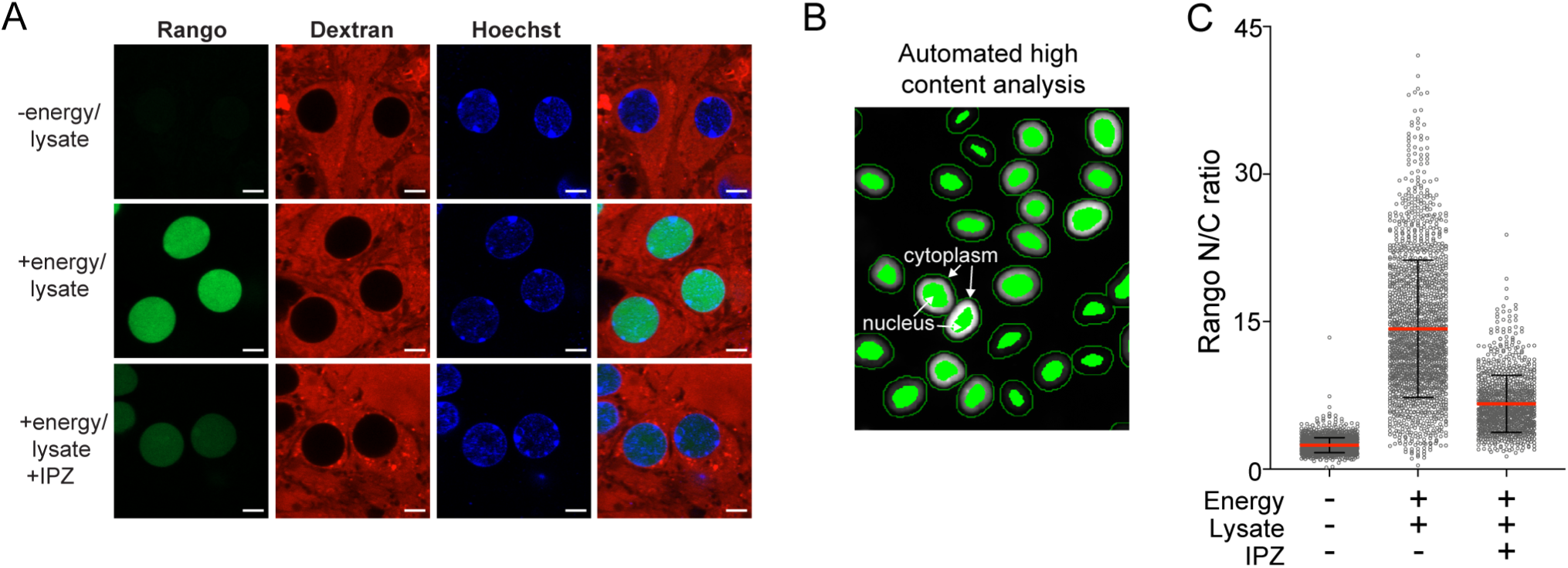
Validation of permeabilized cell assay. **A.** Mouse primary cortical neurons were permeabilized with hypotonic buffer + BSA cushion, and incubated for 2 hours with 200 nM Rango sensor in the indicated conditions. Scale bar = 5µm. **B.** Automated method for N/C ratio calculates mean intensity at a defined distance inside and outside the nuclear rim (as determined by Hoechst signal). **C.** Sample raw data for conditions in **(A),** from a single well of a 96-well plate.

**Figure 1-figure supplement 2.**
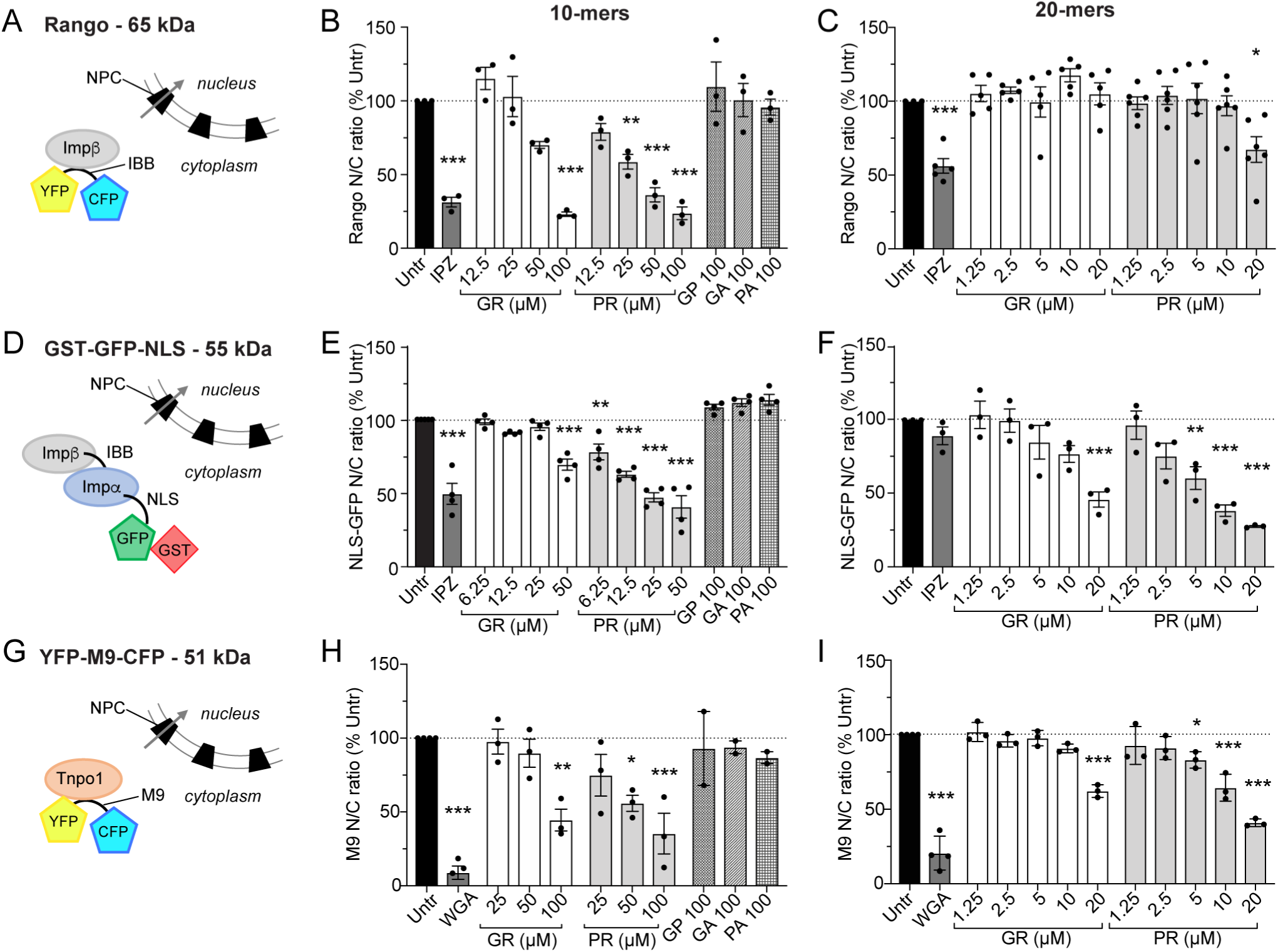
Extended nuclear import data. Diagram of cargo import mechanisms and HeLa nuclear import data in the presence of increasing concentrations of DPR 10- and 20-mers, at steady state (2h for Rango (**A-C),** 4h for NLS-GFP (**D-F**), and 2h for M9 (**G-I**). These data correspond to those summarized in the table in figure 1L. Mean ± SEM is shown, n≥3 biological replicates for R-DPRs, n≥2 biological replicates for GP, GA, and PA. 1209 ± 305 cells per replicate, \**p*<0.05, \*\**p*<0.01, \*\*\**p*<0.001 vs. untreated cells, one-way ANOVA with Dunnett’s post-hoc test. See source file for raw data and exact *p* values.

**Figure 1-source data.** Raw data and *p* values for data in figure 1 and supplements.

**Figure 2-figure supplement 1.**
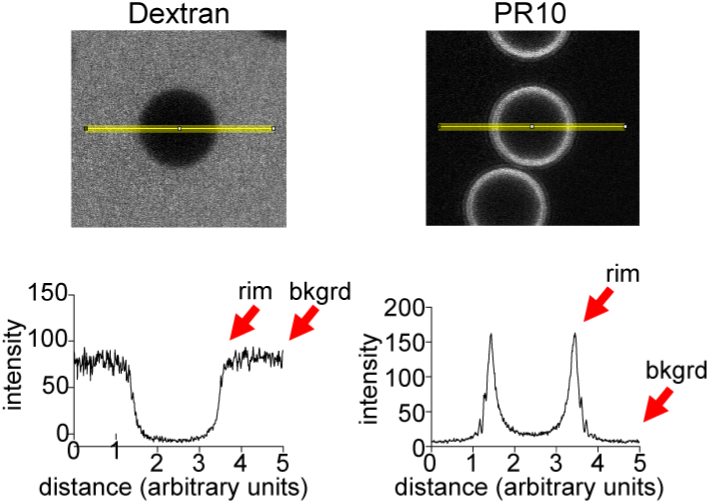
Quantification method for bead halo assay. Examples of line intensity profiles for control versus PR10 beads (Fiji), with rim vs. background levels indicated.

**Figure 2-source data.** Raw data and *p* values for data in figure 2.

**Figure 3-figure supplement 1.**
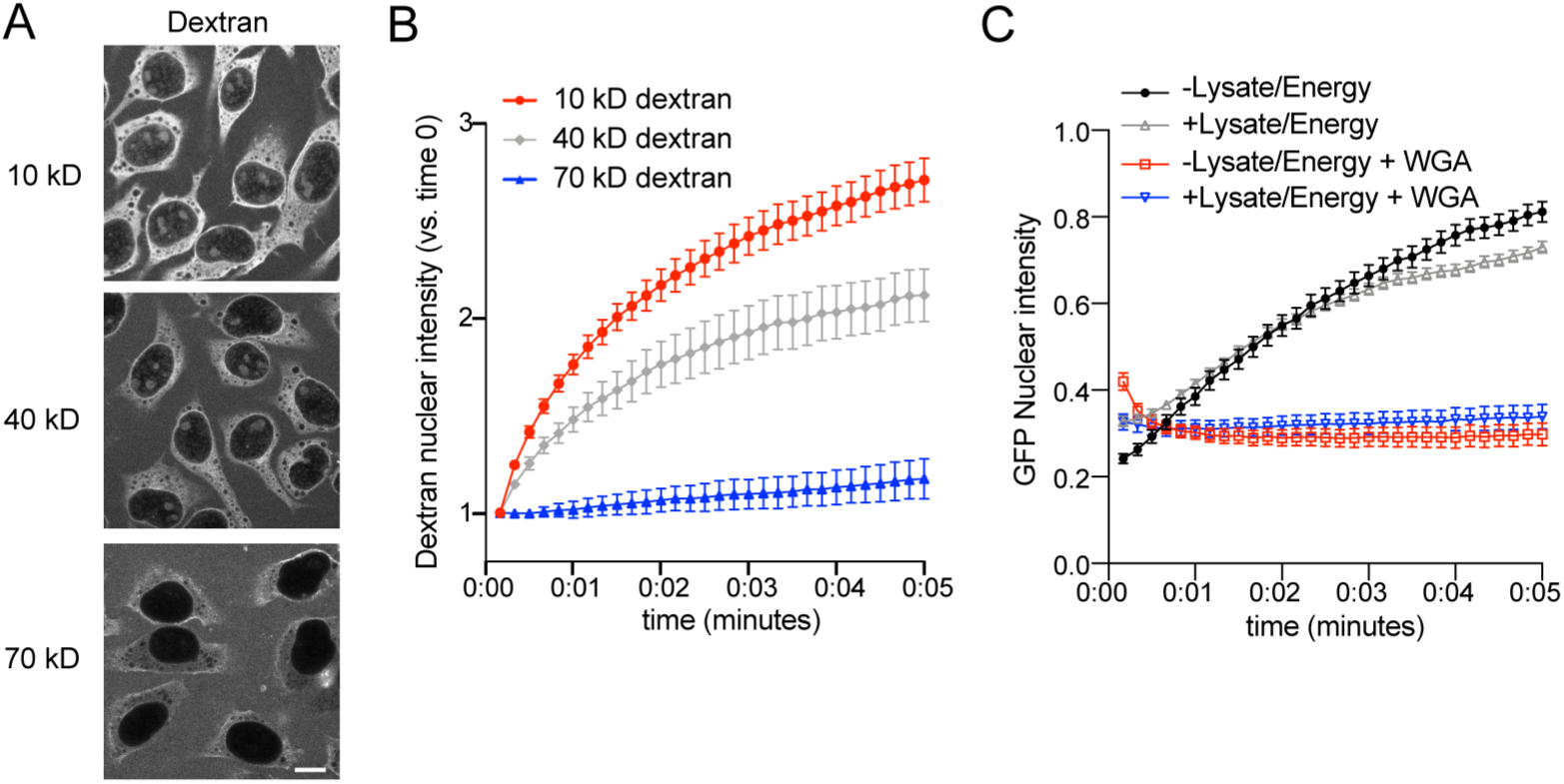
Validation of passive nuclear influx assay. **A.** Confocal images of permeabilized HeLa cells incubated with Texas Red-labeled dextrans of the indicated molecular weight for 15 minutes. Scale bar = 10µm. **B.** Time lapse imaging of dextran nuclear influx from 0-5 minutes. Nuclear intensity is normalized to time 0 for each cell (1=no influx). Mean ± SEM is shown for n=3 biological replicates (20-30 cells/replicate/condition). **C.** Time lapse imaging of GFP nuclear influx, with or without lysate/energy, to verify passive transport of this 27 kD, non-NLS-containing protein. A subset of cells were pre-incubated with 0.8 mg/ml WGA as a positive control for impediment to transport. n=1 (20-30 cells/condition).

**Figure 3-figure supplement 2.**
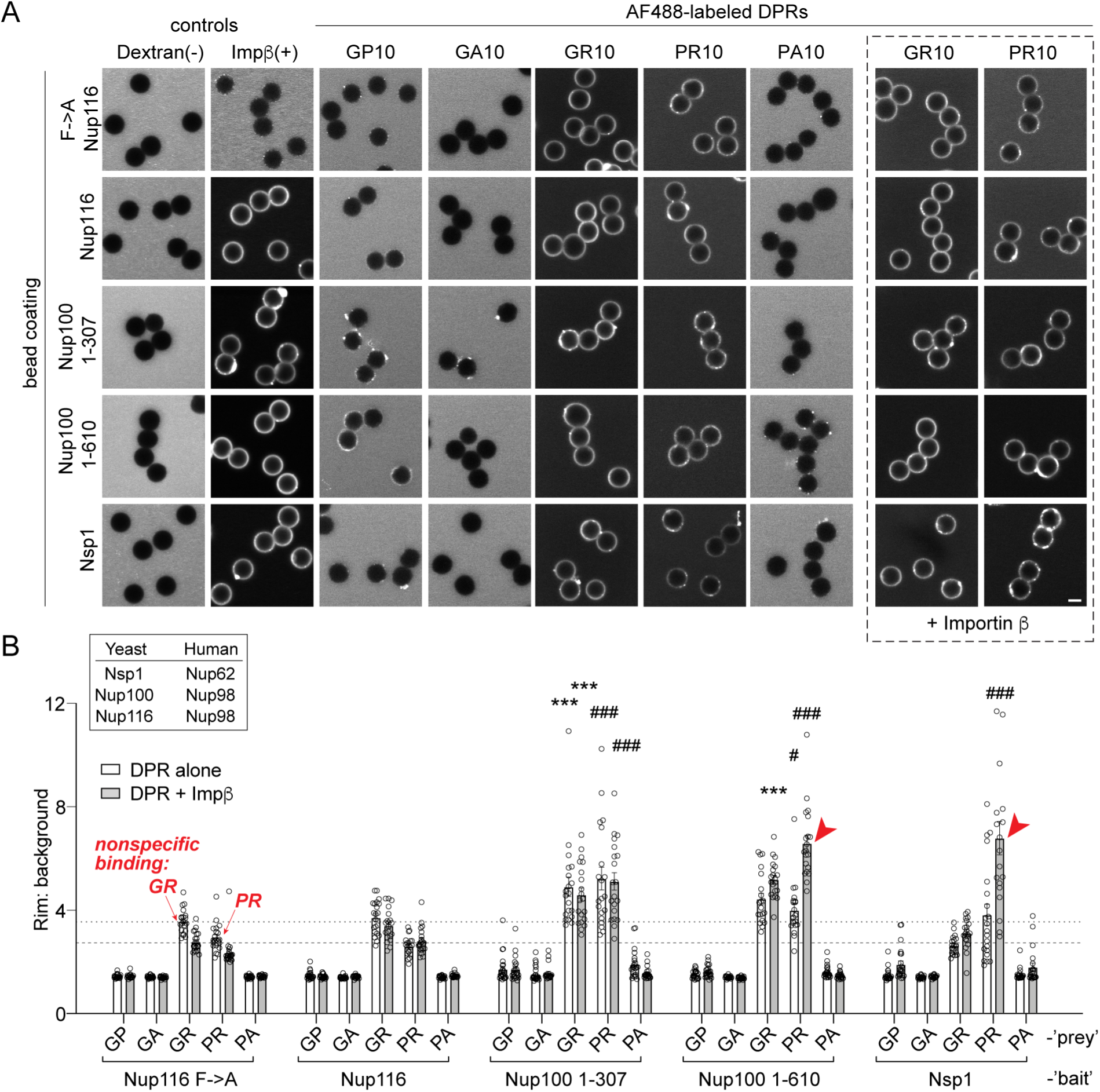
R-DPRs show modest binding to FG-domains in the bead halo assay, which can be augmented by importin β. **A.** Confocal images of AF488-labeled DPRs added to glutathione beads coated with yeast FG- and GLFG-domain GST-fusion proteins, in binding buffer with or without added unlabeled importin β. FITC-dextran = negative control (-), full length AF647-importin β = positive control (+). Scale bar = 4µm. **B.** Intensity profiles (rim vs. background) across all beads tested, including the Nup116 Fà A mutant which is used to define the background/non-specific binding level as indicated by the horizontal dashed lines. Correspondence between yeast and human Nups is given in the inset. Mean ± SEM is shown, for n=20 beads (5 intensity profiles/bead). \**p*<0.05, \*\**p*<0.01, \*\*\**p*<0.001 vs. Nup116 FàA by two-way ANOVA with Tukey post-hoc test (*denotes GR statistics, # denotes PR statistics, red arrows denote augmentation of binding by importin β (*p*<0.001)). See source file for raw data and exact *p* values.

**Figure 3-source data.** Raw data and *p* values for data in figure 3 and supplements.

**Figure 4-figure supplement 1.**
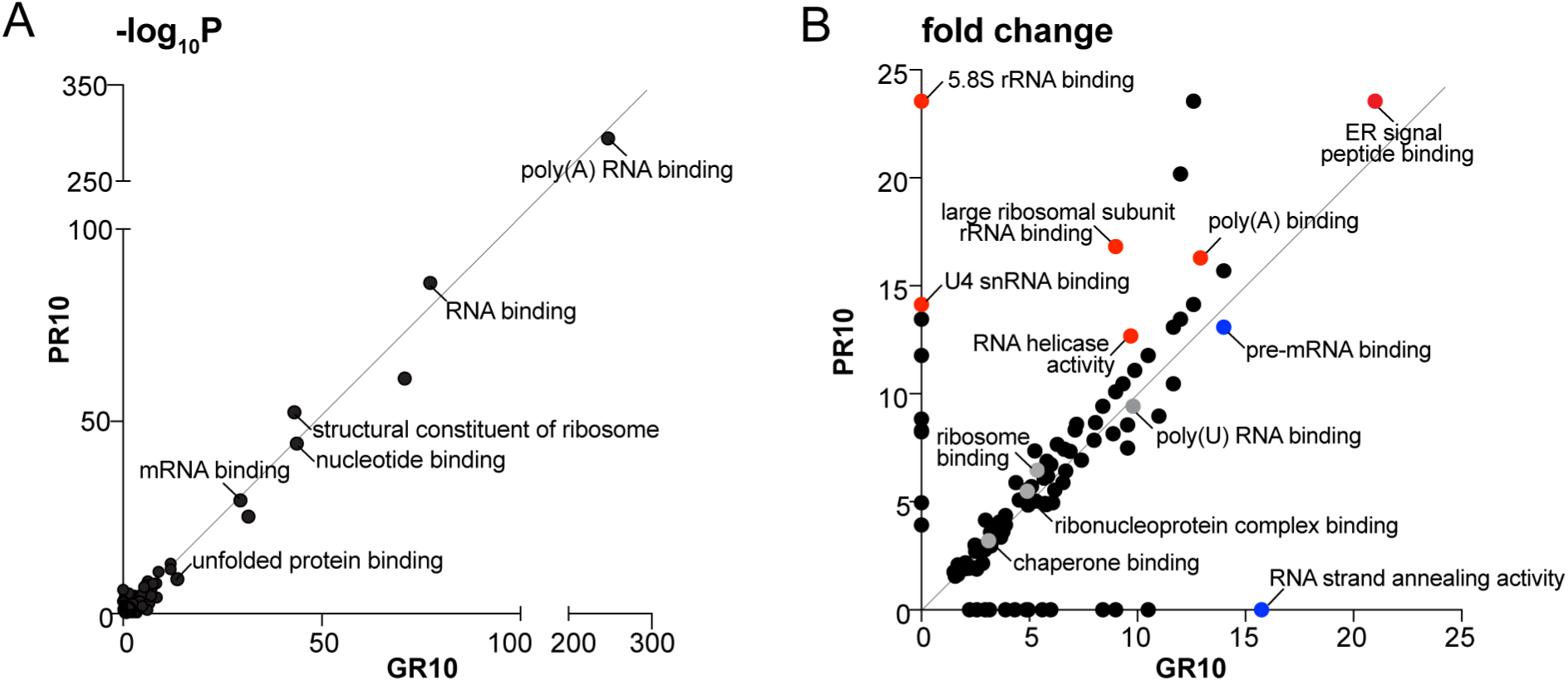
Overall top GO terms enriched in R-DPR aggregates. **A-B.** Top molecular function GO terms for GR10 and PR10 aggregates according to *p* value (shown as −log_10_) (**A**) and fold change (**B**). In B, selected GO categories enriched in PR samples are highlighted in red, and GR in blue.

**Figure 4-figure supplement 2.**
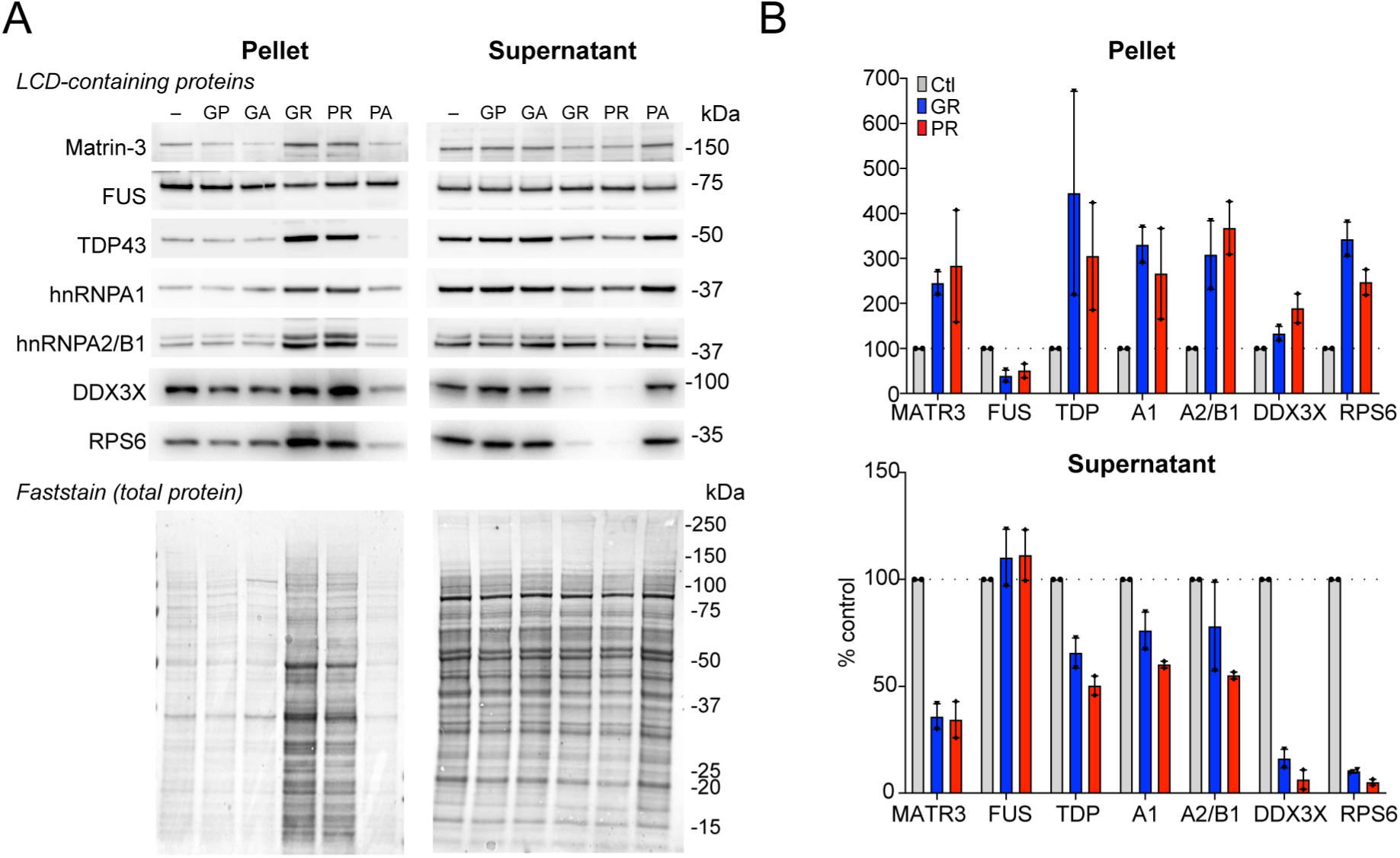
Western blots for selected low complexity-domain (LCD)-containing proteins in R-DPR supernatant vs. pellet fractions. **A.** Western blot for indicated proteins identified by R-DPR aggregate mass spectrometry in supernatant vs. pellet, loaded by volume. A representative post-transfer Faststain (total protein stain) is shown. **B.** Quantification of blots in (**A**). Mean ± SD is shown for two technical replicates. See source file for raw data.

**Figure 4-source data.** Raw data and *p* values for data in figure 4 and supplements.

**Figure 5-figure supplement 1.**
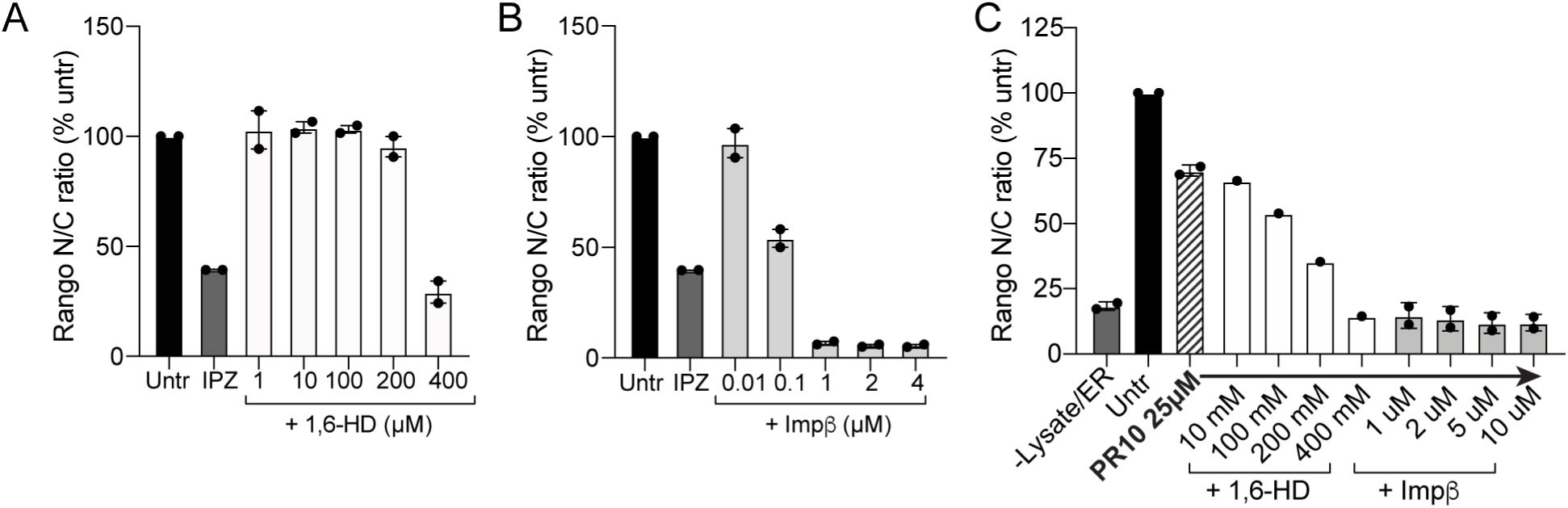
1,6-HD and importin β do not rescue nuclear import in the permeabilized cell assay. **A-B.** 1,6-hexanediol (1,6-HD) (**A**) and WT importin β (**B**) cause dose-dependent inhibition of Rango import in HeLa cells at baseline (mean ± SEM for n=2 replicates is shown). **C.** No rescue of mild Rango import inhibition (25 µM PR10) was seen for either intervention (n=1 for 1,6-HD, and n=2 replicates for importin β, 1622 ± 271 cells/data point). Note that values in C are not background corrected as some fell below the level observed for cells without energy or lysate added. See source file for raw data.

**Figure 5-figure supplement 2.**
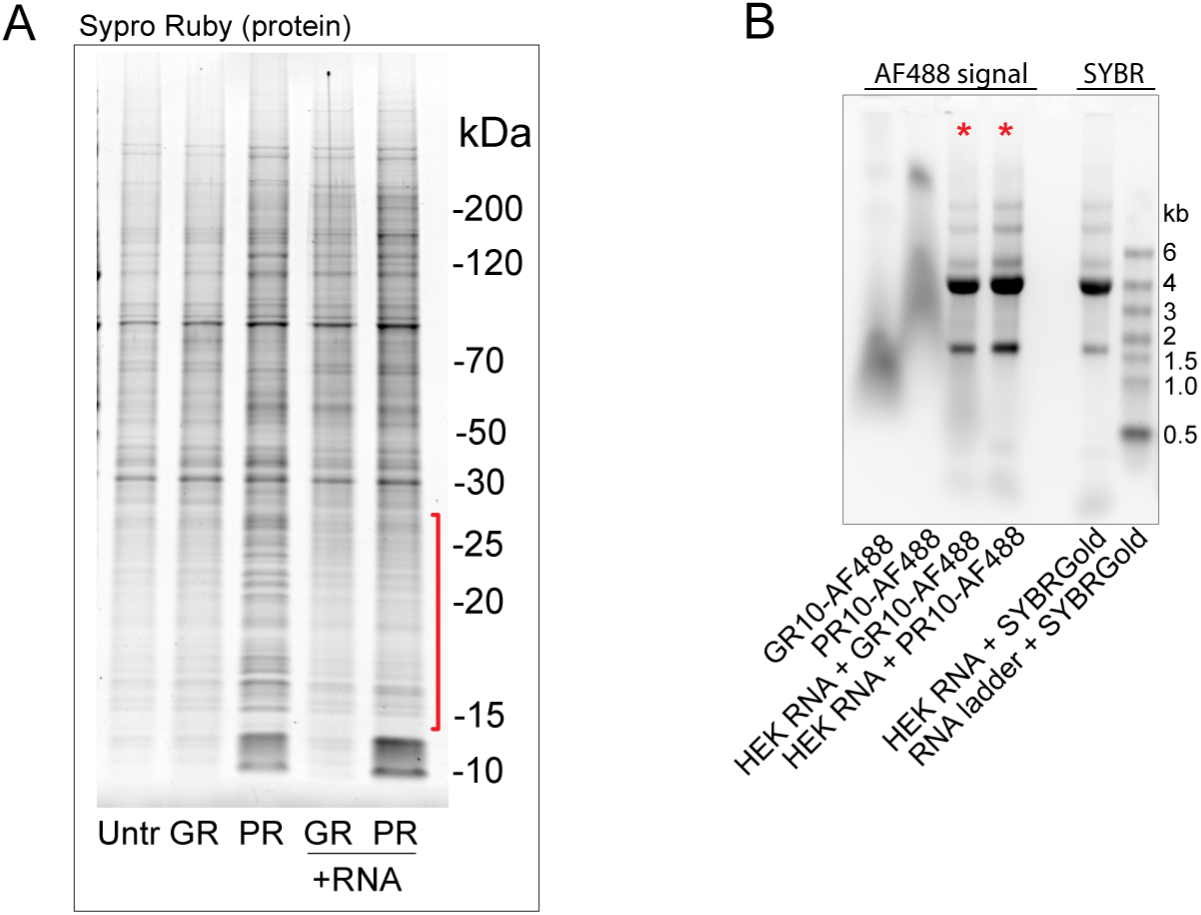
RNA modestly decreases R-DPR aggregates and binds R-DPRs in an electrophoretic mobility shift assay. **A.** Sypro Ruby-stained protein gels showing effect of total HEK RNA on R-DPR-mediated aggregate formation (HEK lysate pellets were prepared as for mass spec and Western blots in figure 4, +/− total HEK RNA). Only a modest reduction of predominantly low molecular weight species was seen in the pellets (bracketed in red). **B.** Electrophoretic mobility shift assay of AF488-labeled DPRs (10mers), +/− total HEK RNA, imaged by UV transillumination to simultaneously visualize the AF488 and SYBR Gold signals. Note the co-migration of AF488 R-DPRs with RNA, as visualized by AF488. No SYBR Gold was added to these lanes (*).

**Figure 5-source data.** Raw data and *p* values for data in figure 5 and supplements.

